# Notch mediates non-regenerative alveolar repair after staphylococcal lung injury

**DOI:** 10.64898/2026.02.04.703850

**Authors:** Sarah K.L. Moore, Stephanie Tang, Deebly Chavez, Sayahi Suthakaran, Chaya Sussman, Jimmy Zhang, Jaime L. Hook

**Author notes:** Corresponding author: Jaime L. Hook, M.D., Icahn School of Medicine at Mount Sinai One Gustave L. Levy Place, Box 1232 New York, N.Y. 10029.

## Abstract

Severe lung infection causes dysfunction of the lung’s air-blood barrier, leading to respiratory failure. In alveoli of lungs infected with *Staphylococcus aureus* (SA) – either alone or after respiratory viral infection – the SA toxin, alpha hemolysin (Hla) causes epithelial barrier protein degradation and airspace edema formation. How the barrier repairs is not clear. We used confocal imaging of intact, perfused, SA-infected lungs to define barrier repair mechanisms in live alveoli. Though we expected to find the non-surviving alveolar epithelium was regenerated, we found, instead, the surviving alveolar epithelium spontaneously regained barrier function. Thus, SA stimulated Notch protein cleavage in the alveolar epithelium in an Hla- and ADAM10-dependent manner. Subsequent exposure of the Notch transmembrane domain catalyzed epithelial junctional protein recovery to reseal the barrier and restore barrier integrity. While disrupting Notch cleavage in the alveolar epithelium prolonged SA-induced lung injury, augmenting it accelerated lung repair. We interpret that barrier repair in the surviving alveolar epithelium resulted from Notch-mediated junctional protein reassembly. These findings show, for the first time, that the alveolar epithelium is a resilient tissue that harbors robust endogenous repair mechanisms. We propose strategies that leverage Notch’s non-regenerative, barrier-strengthening capability may promote lung repair after staphylococcal lung infection.

## INTRODUCTION

When bacteria cause tissue damage, the host must respond by initiating tissue repair pathways that reestablish tissue function. There is little understanding of how bacteria-damaged tissues repair, in part because there is a lack of direct data that inform repair mechanisms in live, intact tissues. This leaves a gap in the understanding of repair mechanisms in tissues that have structural, cellular, and physiological complexity that is difficult to replicate in vitro. Lung alveoli are an example. Alveoli consist of alternating flat and curved septa that separate alveolar airspaces from the fluid-filled lumens of the adjacent capillary meshwork. The intricate alveolar structure includes more than ten epithelial, endothelial, immune, and stromal cell types that are connected by cell-cell communication pathways (1, 2). Still, an accurate understanding of alveolar repair depends on data generated in this complex alveolar microenvironment, since tissue microenvironments determine tissue repair and regeneration mechanisms (3). Here, we aimed to define alveolar repair mechanisms in the microenvironment of live, intact alveoli after lung infection by inhaled *Staphylococcus aureus* (SA).

SA is a common cause of severe lung infection that associates with alveolar damage and high mortality, particularly when it occurs after respiratory viral infection (4–7). The mortality results in part from SA-induced damage to the alveolar barrier, leading to airspace edema formation (8, 9). In health, the alveolar barrier maintains airspace patency by restricting the passage of fluid and proteins from microvascular lumens into airspace lumens. The barrier is composed of alveolar epithelial and microvascular endothelial cells that share a basement membrane (10), and its permeability is regulated by claudin, occludin, and cadherin proteins at alveolar epithelial cell junctions (11–17). SA causes junctional protein degradation in the alveolar epithelium by multiple mechanisms, including interactions between the SA toxin, alpha-hemolysin (Hla) and the epithelial metalloproteinase, ADAM10 that lead to ADAM10-mediated cleavage of epithelial E-cadherin (18). These and other mechanisms lead to barrier hyperpermeability, pulmonary edema formation, and respiratory failure.

The observation that most people survive SA-induced lung injury and regain lung function suggests alveoli have the capacity to repair. How the barrier repairs remains unclear. One possibility is barrier function is restored in the course of alveolar regeneration. If this is the case, then the alveolar epithelium that does not survive SA-induced alveolar damage might be regenerated by alveolar epithelial type 2 (AT2) cells, which proliferate and differentiate into the alveolar epithelial type 1 (AT1) cells that form the bulk of the alveolar surface (19, 20). In this regenerated alveolar epithelium, junctional proteins synthesized de novo might reestablish barrier integrity. An alternative possibility is barrier function is recovered in the alveolar epithelium that survives SA-induced alveolar damage through cellular mechanisms that restore junctional protein expression. Support for this latter possibility comes from cultured endothelial cells, which show robust capacity to spontaneously reassemble junctional proteins and restore endothelial barrier function after exposure to barrier-deteriorating stimuli, such as H_2_O_2_ and thrombin (21–23).

In this report, we outline findings from the first real-time determinations of alveolar barrier repair mechanisms in live, intact, perfused lungs. Our data show alveolar barrier function spontaneously recovered after SA-induced barrier loss. Specifically, SA lung infection stimulated ADAM10- and gamma-secretase-dependent cellular responses in the alveolar epithelium that led to cleavage of the Notch protein. Subsequent exposure of the Notch transmembrane domain (TMD) initiated junctional protein reassembly and recovery of alveolar barrier function. These findings show that, in the alveolar epithelium that survives SA-induced lung injury, the injury stimulates non-regenerative, Notch-dependent mechanisms of tissue repair that recover barrier function. These data raise the possibility, for the first time, that therapeutic strategies that stimulate endogenous alveolar barrier repair pathways could promote lung function recovery after lung infection.

## RESULTS

### The alveolar epithelium rapidly recovers barrier function after SA-induced barrier loss

We began by intranasally instilling mice with PBS or GFP-labeled SA (SA^GFP^) of the clinically-relevant (24, 25) strain, USA300 (**Figure 1A**). We excised the lungs 3 h later, then applied our established methods (26, 27) to perfuse and image the live lungs by confocal microscopy from 4 h to 7 h after instillation (**Figure 1, A-B**). Confocal images show inhaled SA^GFP^ formed microaggregates at structural niches of alveoli, where septa converge (**Figure 1C**), in line with our published data (26, 27). To define alveolar barrier function, we used an established method (26–28) in which we transiently added fluorescein isothiocyanate-labeled dextran (20 kDa; Dextran-20) to the lung perfusate solution (**Figure 1, D-G**). In these experiments, restriction of Dextran-20 fluorescence to microvascular lumens signals alveolar barrier function is intact, while leak of Dextran-20 into airspaces identifies alveoli with barrier hyperpermeability, hence loss of alveolar barrier function (26–28) (**Supplemental Figure 1, A-F**). Our findings show Dextran-20 was confined to microvessels in PBS-instilled lungs (**Figure 1, D and G**), but it leaked from microvessels into alveolar airspace lumens in lungs at 4 h after SA^GFP^ instillation (**Figure 1, E and G**). These data confirm our published findings (26, 27) by showing that SA^GFP^ cause alveolar barrier dysfunction within hours of intranasal SA^GFP^ instillation.

**Figure 1.**
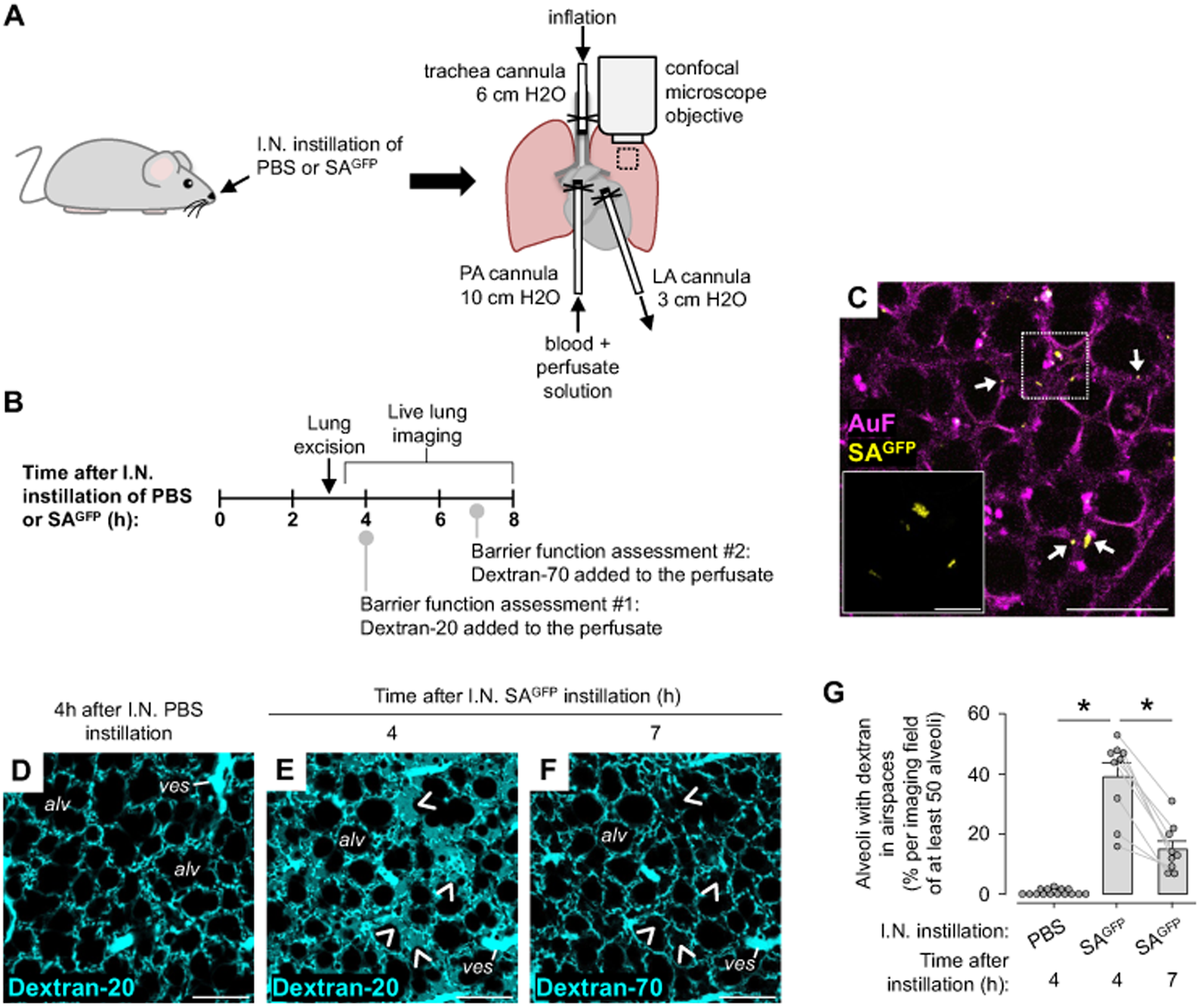
Alveolar barrier function recovers within hours of *S. aureus-*induced barrier loss. **(A-G)** Cartoons (A-B) show the experimental design of imaging studies of alveolar barrier function in live, intact, perfused lungs of mice at the indicated times after intranasal (*I.N.*) instillation of PBS or GFP-tagged *S. aureus* (*SA^GFP^*). Barrier function assessments #1 and #2 were separate determinations of alveolar barrier function that were performed in the same imaging field of at least 50 alveoli by adding fluorophore-tagged dextran to the lung perfusate solution at 4 h (FITC-labeled dextran, 20 kDa, *Dextran-20*) and again at 7 h (TRITC-labeled dextran, 70 kDa, *Dextran-70*) after I.N. instillation. Barrier function was assessed by quantifying dextran leak from microvessels into alveolar airspaces (see Methods). The confocal images in C show low and high (*inset*) power views of SA^GFP^ fluorescence (*yellow*) in airspaces in an example imaging field of alveoli at 4 h after I.N. SA^GFP^ instillation. *Arrows* point out microaggregates of SA^GFP^ at structures niches of alveoli, where septa converge. Autofluorescence (*AuF; magenta*) delineates alveolar walls. *Dashed square* indicates the location of the inset image. Confocal images in D-F show dextran fluorescence (*cyan*) in alveoli. *Arrowheads* indicate locations at which Dextran-20 leaked from microvessels into airspaces during barrier function assessment #1, at 4 h after SA^GFP^ instillation (E), but not during barrier function assessment #2, at 7 h after SA^GFP^ instillation (F). Note, dextran is absent from alveolar airspaces of the PBS-instilled lungs. Group data in G quantify the imaging data in D-F. In G, *gray lines* connect dextran quantifications that were performed at 4 h and 7 h in the same imaging field; circles indicate *n* and each represent one imaging field of at least 50 alveoli in lungs of 3 mice per group; bars: mean ± SEM; \**p* < 0.05 as indicated by ANOVA with post-hoc Tukey testing. Images in C were replicated in 3 mice. Scale bars: 100 μm (C-F) and 20 μm (inset). *PA,* pulmonary artery; *LA*, left atrium; *alv,* example alveolus; *ves,* microvessel.

To define the extent to which alveoli of SA^GFP^-infected lungs recover barrier function, we transiently added fluorophore-labeled dextran to the lung perfusate solution a second time, at 7 h after intranasal SA^GFP^ instillation, and tested alveolar barrier function again in the same alveoli that were imaged at the 4 h time point (**Figure 1B**). For this second determination of barrier function, we added dextran labeled by tetramethylrhodamine isothiocyanate (70 kDa; Dextran-70) to the lung perfusate solution in order to distinguish between dextran that leaked into airspaces at 4 h after instillation from dextran that leaked into airspaces at 7 h after instillation. Confocal images show that, in SA^GFP^-instilled lungs, fewer than half of the alveoli in which Dextran-20 leaked into airspaces at 4 h after instillation now had Dextran-70 leak at 7 h after instillation (**Figure 1, F-G**). We conclude that alveolar barrier function recovered in SA^GFP^-infected lungs. Separate experiments, in which we gave Dextran-20 at both 4 h and 7 h after instillation, show the same findings (**Supplemental Figure 1, G-I**), ruling out the possibility that differences in barrier leak at the two time points resulted from differences in alveolar sieving properties to dextrans of different sizes. Control experiments show that lungs instilled with PBS retained alveolar barrier function at 7 h after instillation (**Supplemental Figure 1, G and J-K**), indicating the live lung imaging procedure did not affect barrier function. These findings provide the first direct evidence, of which we are aware, that alveoli spontaneously regain barrier function within hours of SA^GFP^-induced barrier loss.

To support the live lung imaging data, we tested the extent to which alveoli regain barrier function in a mouse model of lung infection. We intranasally-instilled mice with PBS or SA^GFP^ (**Figure 2A**). All mice survived for at least 48 h after instillation (**Figure 2B**). Quantifications of mouse breathing score, an observational scoring system we developed that correlates with markers of pulmonary edema in mice (27), show that intranasal instillation of SA^GFP^, but not PBS, induced breathing abnormalities within 4 h of instillation (**Figure 2C**). The breathing abnormalities persisted for hours, then normalized by 24 h after SA^GFP^ instillation (**Figure 2C**). These findings show mice rapidly recovered from SA^GFP^-induced lung injury and suggest that SA^GFP^-induced pulmonary edema formed, then resolved over a period of hours.

**Figure 2.**
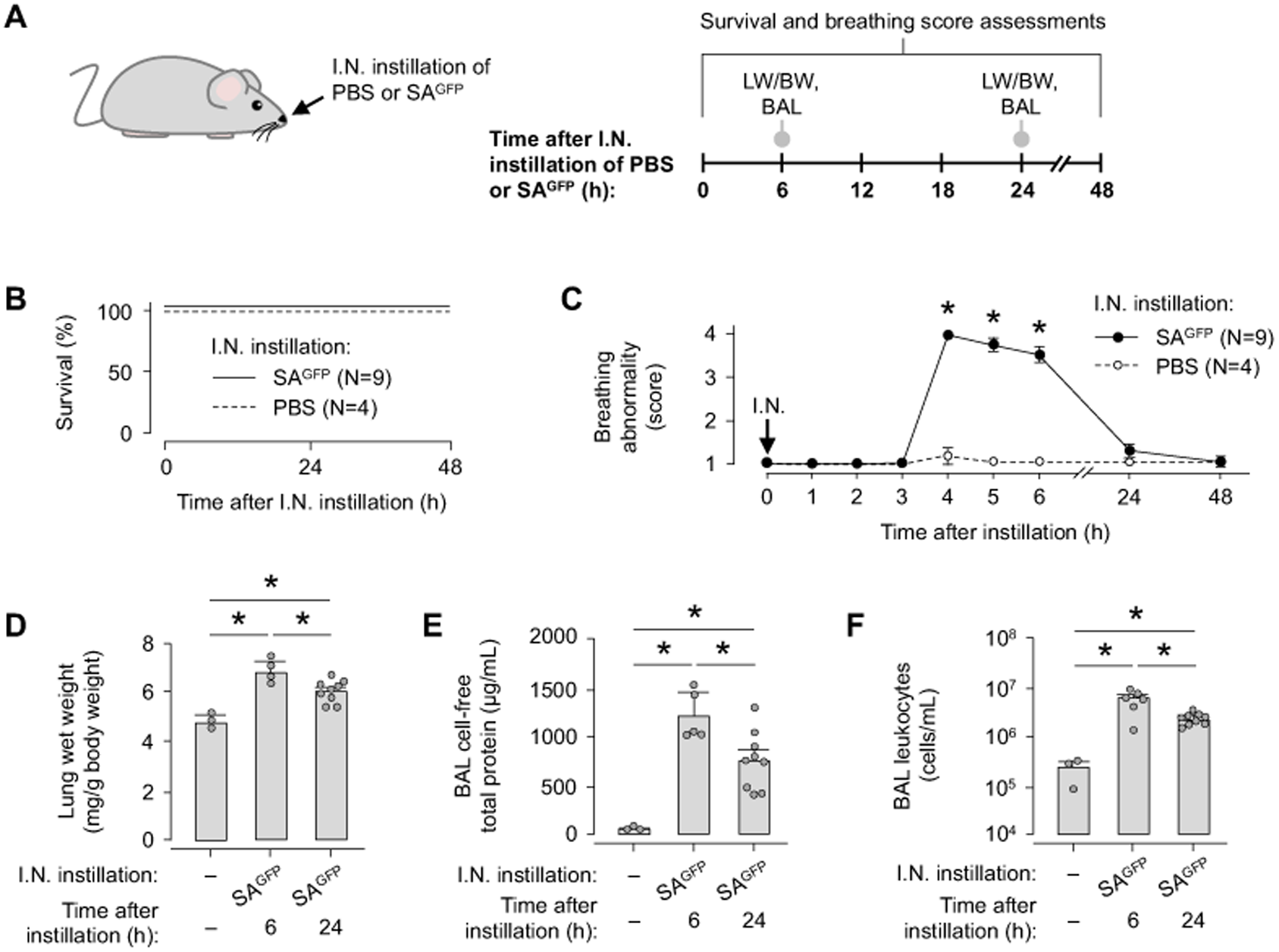
Pulmonary edema resolves after *S. aureus*-induced acute lung injury. **(A-F)** The cartoon in A shows the experimental design of studies used to generate the group data in B-F. As indicated in A, we carried out intranasal (*I.N.*) instillation of PBS or SA^GFP^ in mice, then quantified mouse survival (B), breathing score (C), lung wet weight to body weight ratio (*LW/BW*) (D), and bronchoalveolar lavage (*BAL*) fluid content of protein (E) and leukocytes (F). In B and C, *n* is indicated in the panel legends. In B, lines are not significantly different by log rank test. In C, *circles* indicate mean ± SEM; \**p* < 0.05 versus PBS by two-tailed *t* test. In D-F, *circles* indicate *n* and each represent one mouse; bars: mean ± SEM; \**p* < 0.05 as indicated by ANOVA with post-hoc Tukey testing. BAL fluid contents of protein and leukocytes in E-F were quantified using the same fluid specimens.

Follow up experiments affirm that the breathing changes we observed indeed reflected pulmonary edema formation, then resolution. We quantified two measures of pulmonary edema: lung wet weight to body weight (LW/BW) ratio and bronchoalveolar lavage (BAL) fluid total protein content. Our findings using both measures show lung content of edema fluid increased at 6 h after SA^GFP^ instillation, then decreased at 24 h after instillation (**Figure 2, D-E**). These findings are supported by data that show lung inflammation increased, then decreased within the same time period (**Figure 2F**), bolstering the notion that SA^GFP^-induced lung injury rapidly resolved. Taking these findings together, we conclude that pulmonary edema formed, then resolved within hours of SA^GFP^ instillation. Although the rapid resolution of pulmonary edema probably resulted from a combination of alveolar barrier repair and alveolar fluid clearance, the imaging data in **Figure 1, E-G** show that barrier repair was a critical factor in the edema resolution.

### The alveolar epithelium retains viability in SA-infected lungs

Next, we aimed to define mechanisms by which the alveolar barrier repaired. We considered that the repair could result from either the de novo establishment of barrier function in alveoli undergoing alveolar epithelial regeneration or from the recovery of barrier function in the surviving alveolar epithelium. Since our published data show the alveolar epithelium retains viability after plasma membrane damage by the pore-forming SA toxin, Hla (26), we tested the hypothesis that the alveolar epithelium that has barrier hyperpermeability survives SA lung infection (**Figure 3, A-E**) and can therefore undergo non-regenerative repair. To do this, we instilled alveolar airspaces of SA^GFP^-infected lungs with the cell-permeant viability dye, calcein red-orange AM, as we have done previously (26, 27). Confocal images show alveoli of SA^GFP^-instilled lungs are visible using the high laser power and gain required to view autofluorescence, but not at lower levels of power and gain needed to view calcein fluorescence after epithelial loading (**Figure 3, A-B**). However, calcein microinstillation in the same alveoli led to robust epithelial fluorescence in lungs at 4 h after intranasal SA^GFP^ instillation (**Figure 3C**), indicating that the alveolar epithelium retained viability at 4 h after SA^GFP^ lung infection. Calcein fluorescence decreased over time, consistent with our published data (26) (**Figure 3D**), but it again increased after repeat calcein microinstillation at 7 h after intranasal SA^GFP^ instillation (**Figure 3E**). By contrast, calcein fluorescence was absent in the epithelium of alveoli treated with airspace instillation of saponin at 1% w/v, which rapidly causes cell death (29) (**Figure 3, F-G**). Taking the group data together (**Figure 3H**), we conclude that the alveolar epithelium of SA^GFP^-infected lungs retained viability at both 4 h and 7 h after SA^GFP^instillation. Thus, we interpret that the alveolar barrier repair we observed within hours of SA^GFP^ lung infection resulted from rapid recovery of barrier function in the surviving alveolar epithelium, rather than de novo establishment of barrier function after alveolar regeneration – a process that takes days to weeks to occur (30).

**Figure 3.**
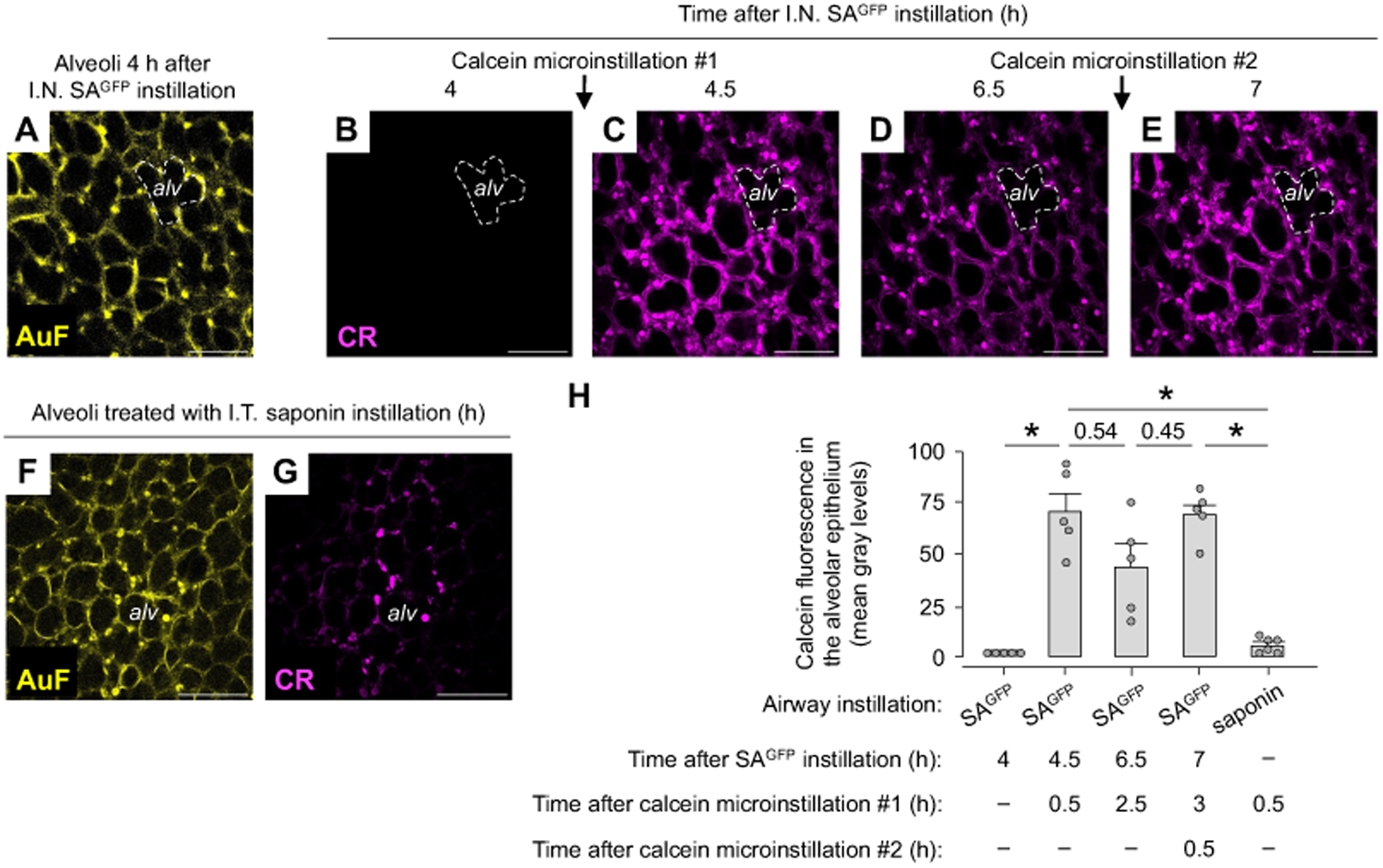
The alveolar epithelium retains viability despite *S. aureus*-induced barrier loss. **(A-H)** Confocal images in A-E show an example imaging field of approximately 50 alveoli at the indicated times after intranasal (*I.N.*) instillation of SA^GFP^. Images were generated before (A-B) and after (C-E) the alveolar airspaces were microinstilled with the viability dye, calcein red-orange AM (*CR; magenta*). A and B are the same image but at imaging settings optimized for alveolar autofluorescence (*AuF*; *yellow*; A) and calcein (B). Images in B-E were generated using the same imaging settings. *Dashed line* in A-E outlines an example alveolus. Confocal images in F-G show an example imaging field of approximately 50 alveoli that were treated with intratracheal (*I.T.*) instillation of saponin prior to alveolar calcein microinstillation. Note, calcein fluorescence in the alveolar epithelium is detectable after calcein microinstillation in alveoli of SA^GFP^-instilled lungs (C-E), but not saponin-instilled lungs (G). Group data in H quantify the imaging findings. In H, circles indicate *n* and each represent one imaging field of at least 50 alveoli; bars: mean ± SEM; \**p* < 0.05 or as indicated by ANOVA with post-hoc Tukey testing. All images were replicated in the lungs of at least 2 mice per group. Scale bars: 100 μm. *alv,* example alveolus.

### SA induces Notch cleavage in the alveolar epithelium

Since reports indicate that cellular pathways involving Notch cleavage are barrier-strengthening in cultured endothelial cells (31), we considered that Notch cleavage might have a role in alveolar barrier repair in SA^GFP^-infected lungs. This possibility is supported by reports that identify Notch expression in the adult alveolar epithelium (32) and Notch cleavage in endothelial cells exposed to Hla (33). To define whether Notch cleavage has a role in alveolar barrier repair, we first aimed to identify if SA^GFP^ instillation causes Notch cleavage in the alveolar epithelium. To do this, we intranasally-instilled mice with 12XCSL-DsRed-ExpressDL, a plasmid that signals Notch cleavage by generating DsRed fluorescence that initiates within 2 h of Notch receptor activation and persists for longer than 8 h (34). We and others have shown previously that intranasal instillation of liposome-complexed plasmid DNA leads to plasmid expression in lung alveoli (26, 27, 35).

To define Notch responses to SA lung infection, we intranasally-instilled mice with the 12XCSL-DsRed-ExpressDL Notch cleavage reporter to generate reporter expression in the alveolar epithelium. To test the hypothesis that SA causes Notch cleavage, we followed the DsRed-reporter plasmid instillation with intranasal instillation of PBS or SA (**Figure 4, A-E**). In lungs of mice instilled with the DsRed-reporter and PBS, confocal images showed weak DsRed fluorescence in the alveolar epithelium (**Figure 4, B and E**). DsRed-reporter fluorescence was nearly doubled in the alveolar epithelium of mice instilled with the DsRed-reporter plasmid and SA^GFP^ (**Figure 4, C and E**), indicating that SA^GFP^ instillation caused Notch cleavage in the alveolar epithelium. Since Hla causes Notch cleavage in cultured endothelial cells (33), we tested the hypothesis that Hla is required for Notch cleavage in alveoli of SA-infected lungs. In line with this hypothesis, we found no increase of DsRed-reporter fluorescence in mice that were intranasally instilled with a GFP-expressing, mutant USA300 SA that is Hla-deficient (**Figure 4, D-E and Supplemental Figure 2, A-B**), indicating that SA^GFP^-induced Notch cleavage is Hla-dependent. We validated the assumption that the SA^GFP^-induced increase of DsRed-reporter fluorescence reflected Notch cleavage by quantifying SA^GFP^-induced DsRed-reporter fluorescence changes in mice pretreated with intranasal instillation of siRNA (**Figure 4, F-J**). siRNA was non-targeting or against presenilin1 (siPS1), the catalytic subunit of the sole Notch-cleaving enzyme, gamma-secretase (36). siPS1 blocked the SA^GFP^-induced increase of DsRed-reporter fluorescence (**Figure 4, F-H and J**), affirming that the increase of DsRed-reporter fluorescence in SA^GFP^-instilled lungs resulted from Notch cleavage. Finally, since Notch is cleaved by ADAM10 (37) and ADAM10 is activated by Hla (18), we tested the hypothesis that ADAM10 is required for Notch cleavage in alveoli of SA^GFP^-infected lungs. SA^GFP^ instillation failed to cause DsRed-reporter fluorescence to increase in mice pretreated with siRNA against ADAM10 (**Figure 4, F and I-J**), indicating that SA^GFP^-induced Notch cleavage was ADAM10-dependent. Immunoblots of cultured murine alveolar epithelial-like cells support the live lung imaging data by affirming that Hla exposure caused Notch1 cleavage (**Figure 4, K-L**) and Hes1 transcription (**Figure 4, M-N**), a marker of Notch cleavage and subsequent release of the transcriptional activator, the Notch intracellular domain (NICD). Taking these findings together, we conclude that SA^GFP^ lung infection caused Notch1 cleavage in the alveolar epithelium through Hla-epithelial interactions and epithelial ADAM10 activation.

**Figure 4.**
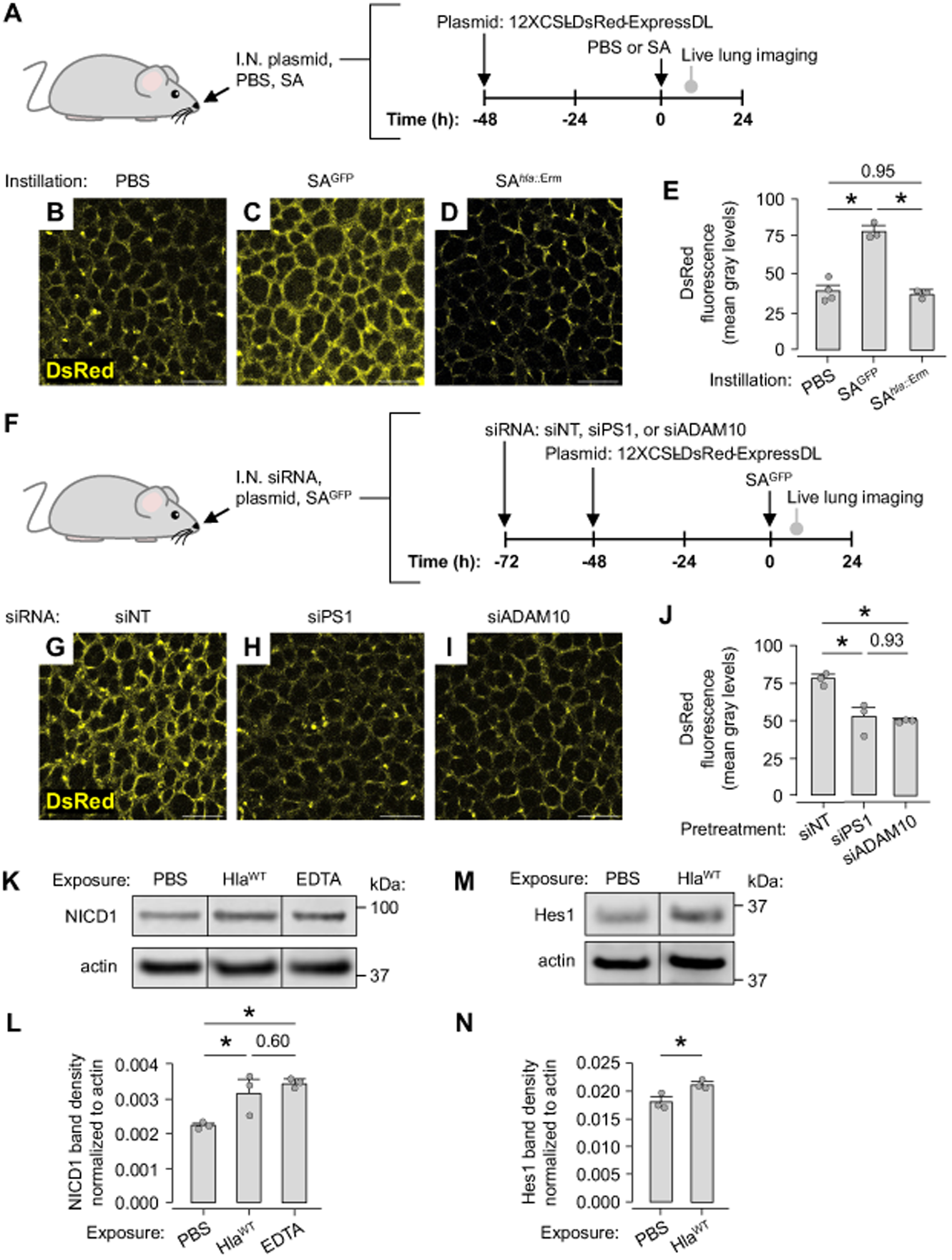
*S. aureus* stimulates Notch cleavage in the alveolar epithelium. **(A-J)** Cartoons in A and F show the experimental design of the studies used to generate the data shown in B-E and G-J, respectively. As indicated in A and F, we gave intranasal (*I.N.*) instillation of the 12XCSL-DsRed-ExpressDL Notch cleavage reporter plasmid, then I.N. instillation of PBS or *S. aureus* (SA). SA were SA^GFP^ or SA*^hla::^*^Erm^ as indicated. At 4 h after PBS or SA instillation, we excised the lungs and used confocal microscopy to view DsRed fluorescence (*yellow*) in alveoli. As indicated in F, mice used to generate the data shown in G-J were also intranasally-instilled with siRNA that was non-targeting (*siNT*) or against presenilin1 (*siPS1*) or ADAM10 (*siADAM10*). In E and J, *circles* indicate *n* and each represent one mouse in which mean DsRed fluorescence was quantified in imaging fields of at least 50 alveoli; bars: mean ± SEM; \**p* < 0.05 or as indicated by ANOVA with post-hoc Tukey testing. Scale bars: 100 μm. (**K-N**) Immunoblots (K and M) and group band densitometry (L and N) show protein content in cultured alveolar epithelial-like MLE-12 cells. Prior to immunoblot processing, the cells were exposed to PBS, purified Hla (*Hla^WT^*), or EDTA as indicated for 10 min, then incubated in culture media for 45 min. EDTA served as a positive control. Bands shown in K and M were generated on the same blot but are not contiguous. In L and N, bars show mean ± SEM; \**p* < 0.05 or as indicated by ANOVA with post-hoc Tukey testing (L) or two-tailed *t* test (N); *circles* indicate *n* and each represent one biological replicate. Replicates were run in the same gel and immunoblotted on the same membrane. Immunoblots were each repeated 2x with the same results.

To rule out the possibility that reporter plasmid expression was in the microvascular endothelium, we carried out a series of experiments in lungs treated with intranasal instillation of a plasmid that encodes GFP (**Figure 5, A-D**). First, we identified by confocal microscopy that plasmid instillation led to widespread GFP fluorescence in alveoli (**Figure 5A**), as expected. Next, we transiently added to the perfusate either vehicle solution or vehicle solution containing the GFP fluorescence-quenching agent, copper sulfate (CuSO_4_). GFP fluorescence did not change (**Figure 5, B and D**), suggesting GFP expression was not in the endothelium. However, CuSO_4_ microinstillation into airspaces caused major GFP fluorescence loss (**Figure 5, C-D**). Since plasmid fluorescence was quenched by copper exposure from airspace lumens but not vascular lumens, we conclude that intranasal plasmid instillation leads to plasmid expression in the alveolar epithelium, but not the microvascular endothelium. These findings strengthen our conclusion that Notch cleavage was localized to the alveolar epithelium of SA^GFP^-infected lungs.

**Figure 5.**
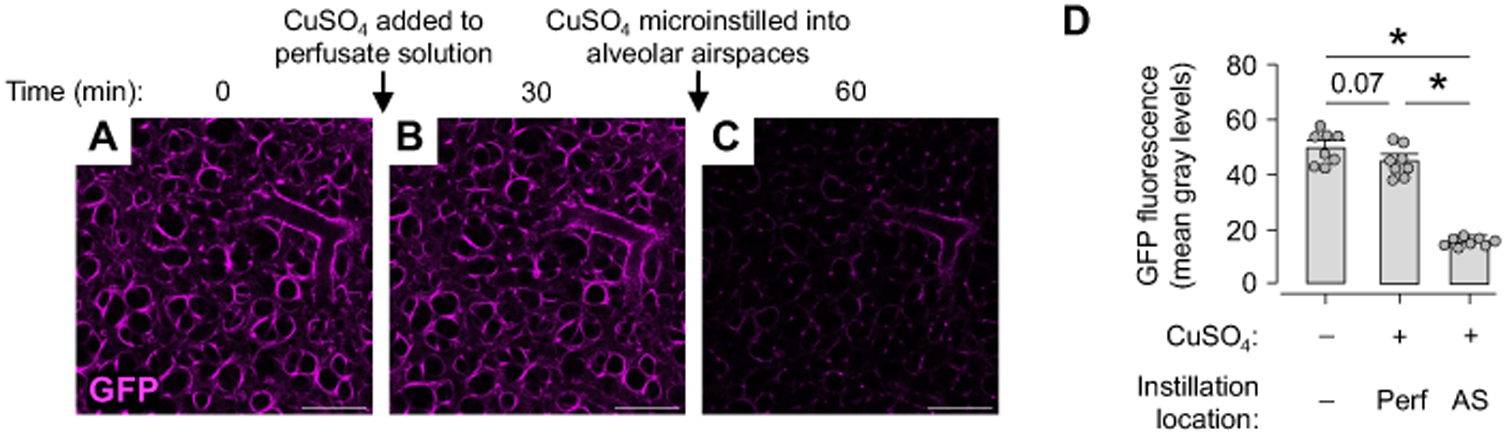
Intranasal plasmid instillation leads to plasmid expression in the alveolar epithelium. **(A-D)** Confocal images in A-C show GFP fluorescence (*magenta*) in an imaging field of approximately 50 alveoli in the lungs of a mouse that was pretreated with intranasal instillation of liposome-complexed, GFP-encoding plasmid DNA. Note, GFP fluorescence persists after lung perfusion with the GFP-quenching agent, CuSO_4_, but it is lost after microinstillation of CuSO_4_ into alveolar airspaces. Group data in D quantify the imaging findings. In D, circles indicate *n* and each represent one imaging field of at least 50 alveoli; bars: mean ± SEM; \**p* < 0.05 or as indicated by ANOVA with post-hoc Tukey testing. Scale bars: 100 μm.

### Notch cleavage mediates alveolar barrier repair after SA lung infection

We next defined whether Notch cleavage in the alveolar epithelium mediates alveolar barrier repair. To do this, we applied the same methods we used previously (**Figure 1**) to intranasally-instill mice with SA^GFP^, then excise the lungs for live lung imaging at 3 h after SA^GFP^ instillation. We again identified alveoli with barrier dysfunction by transiently adding Dextran-20 to the lung perfusate solution at 4 h after SA^GFP^ instillation, then adding Dextran-70 to the perfusate solution at 7 h after instillation (**Figure 6A**). However, at 5 h after instillation, we also added to the lung perfusate solution either DMSO or DMSO in solution with the gamma-secretase inhibitor, DAPT (38) (**Figure 6, A-F**), on the basis that DMSO and DAPT would reach the alveolar epithelium due to alveolar barrier dysfunction or by diffusion, since lipophilic small molecules like DAPT are taken up by the lung after intravenous administration (39). DMSO and DAPT were maintained in the perfusate solution for the remainder of the experiment.

**Figure 6.**
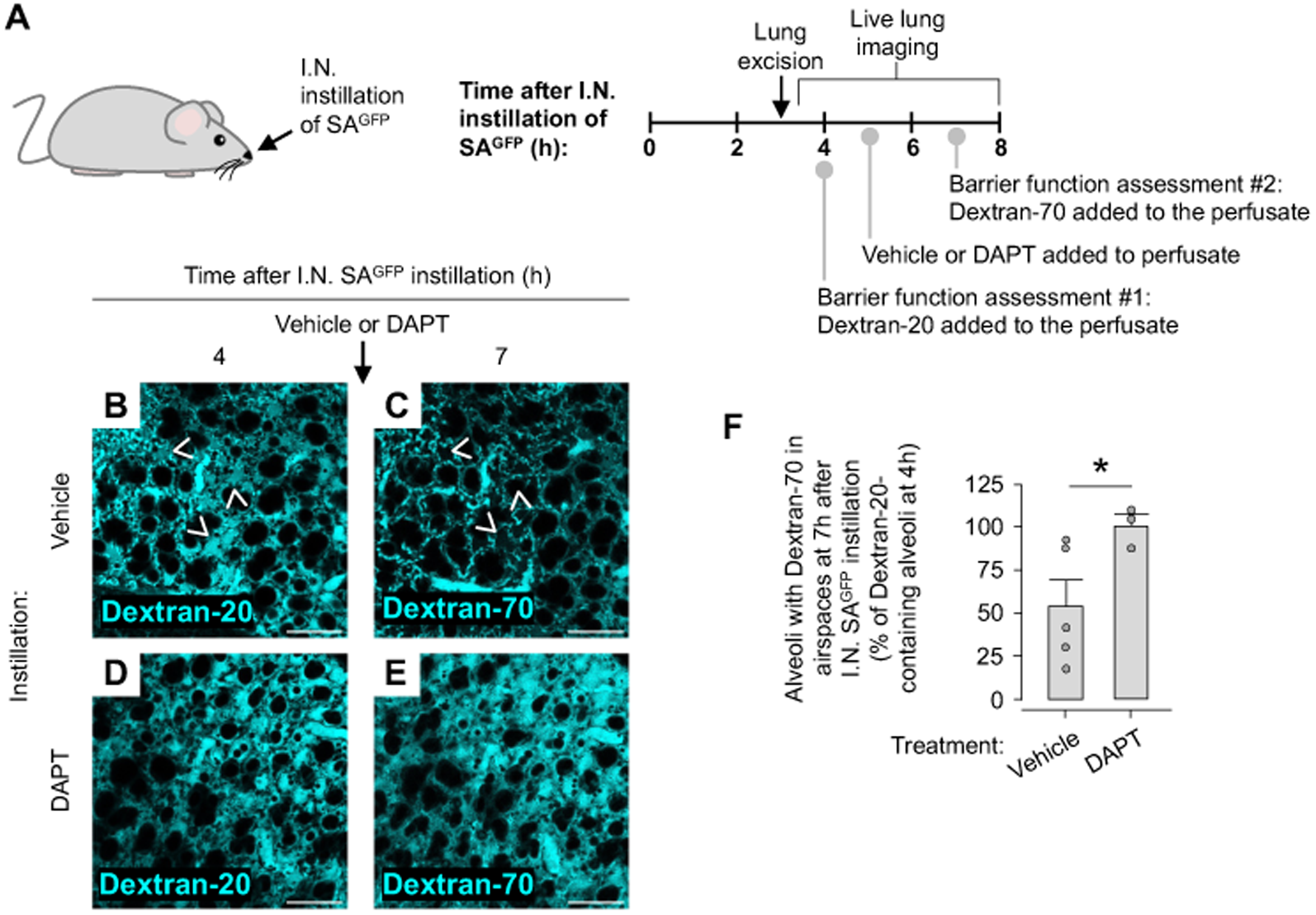
Notch cleavage in the alveolar epithelium mediates alveolar barrier repair. **(A-F)** The cartoon (A) shows the experimental design of studies used to generate the data shown in B-F. As indicated in A, we gave intranasal (*I.N.*) instillation of SA^GFP^, then excised the lungs and viewed the live alveoli by confocal microscopy (B-F). We carried out a three-step experiment in which we: (1) assessed alveolar barrier function at 4 h after SA^GFP^ instillation by adding FITC-labeled dextran (20 kDa, *Dextran-20*) to the lung perfusate solution; (2) added either vehicle or the gamma-secretase inhibitor, DAPT to the lung perfusate solution at 5 h after SA^GFP^ instillation; and (3) reassessed alveolar barrier function at 7 h after SA^GFP^ instillation by adding TRITC-labeled dextran (70 kDa, *Dextran-70*) to the lung perfusate solution. Barrier function was assessed in confocal images (B-E) and associated group data (F) by quantifying dextran leak from microvessels into alveolar airspaces. In images of alveoli exposed to vehicle (B-C), *arrowheads* point out locations at which dextran leaked from microvessels into airspaces at 4 h after SA^GFP^ instillation (B), but not at 7 h after SA^GFP^ instillation (C). Note, dextran fills airspaces of DAPT-exposed alveoli at both time points (D-E). In F, circles indicate *n* and each represent one imaging field of at least 50 alveoli; bars: mean ± SEM; \**p* < 0.05 as indicated by ANOVA with post-hoc Tukey testing. Scale bars: 100 μm.

Our findings show that, as expected, Dextran-20 leaked into alveolar airspaces of SA^GFP^-instilled lungs at 4 h after instillation (**Figure 6, B and D**), indicating that SA^GFP^ caused alveolar barrier loss. Also as expected, at 7 h after instillation, alveoli exposed to vehicle had major reduction of barrier leak (**Figure 6, C and F**), signaling barrier repair. By contrast, alveoli exposed to DAPT had no reduction of barrier leak (**Figure 6, E-F**). We conclude that DAPT blocked the resolution of alveolar barrier dysfunction at 7 h after SA^GFP^ instillation, and we interpret that Notch cleavage mediates barrier repair in SA^GFP^-infected lungs.

We followed up by testing the role of Notch cleavage in barrier repair in mouse models of SA^GFP^ lung infection (**Figure 7A**). Mouse pretreatment with intranasal instillation of siPS1 had no effect on SA^GFP^-induced changes in mouse breathing patterns, LW/BW ratio, or BAL protein content at 6 h after SA^GFP^ instillation (**Figure 7, B-D**), indicating that Notch cleavage in the alveolar epithelium did not determine the extent of the initial SA^GFP^-induced lung injury. However, siPS1 blocked the normalization of the breathing abnormalities, LW/BW ratio, and BAL protein content at 24 h after SA^GFP^ instillation (**Figure 7, B-D**), indicating that inhibition of Notch cleavage in the alveolar epithelium blocked pulmonary edema resolution and disrupted lung repair. Since siPS1 had no effect on BAL leukocyte content or lung SA^GFP^ content at either 6 h or 24 h after SA^GFP^instillation (**Figure 7, E-F**), we rule out the possibility that the injury-prolonging effect of siPS1 was mediated by effects on lung inflammation or bacterial killing. We conclude that Notch cleavage in the alveolar epithelium is required for pulmonary edema resolution after SA^GFP^ lung infection. These mouse model data corroborate the imaging findings (**Figure 6, B-F**) and provide supporting evidence that Notch cleavage in the alveolar epithelium is central to alveolar barrier repair after SA^GFP^-induced lung injury.

**Figure 7.**
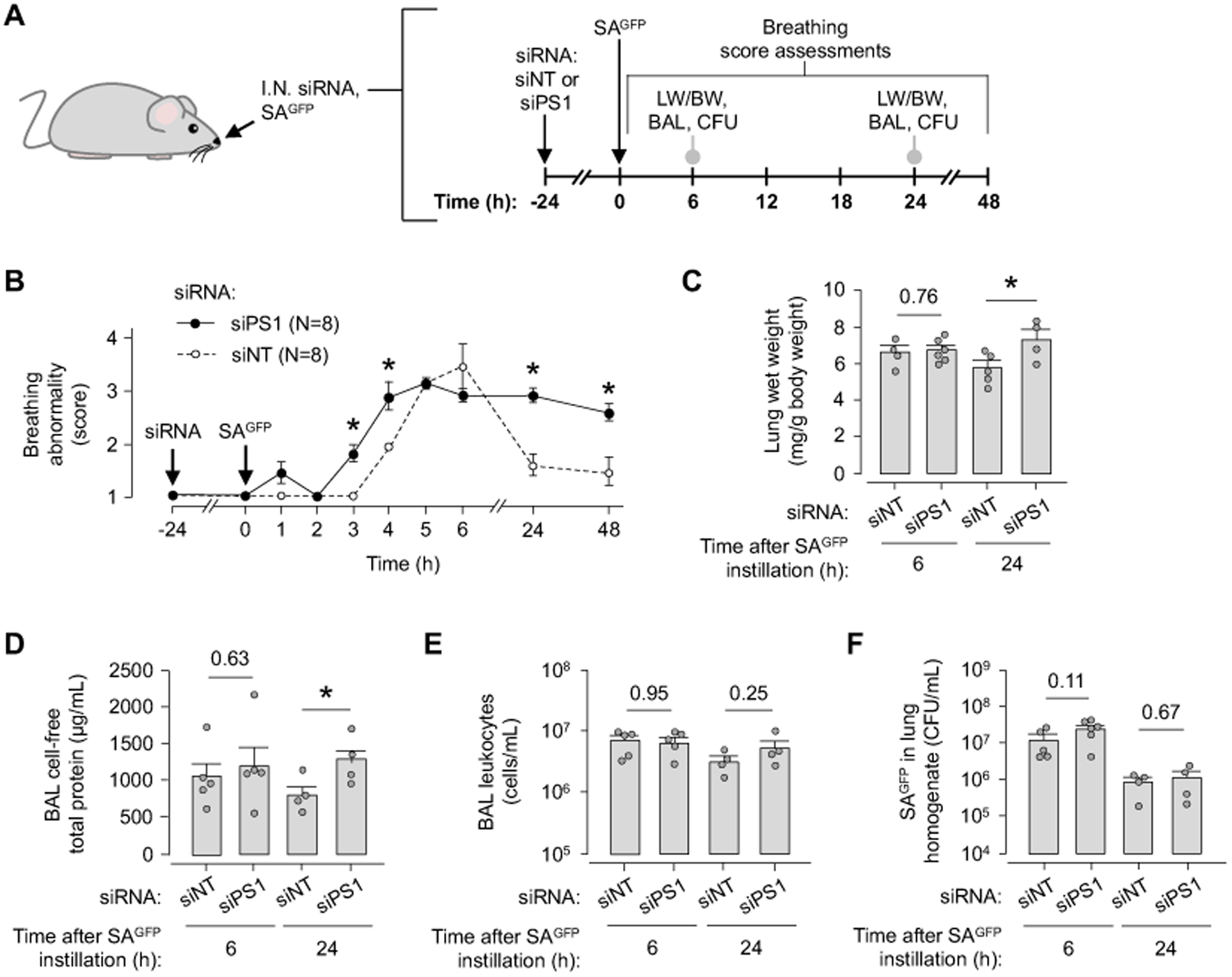
Notch cleavage in the alveolar epithelium promotes lung repair after *S. aureus-*induced acute lung injury. **(A-F)** The cartoon in A shows the experimental design of studies used to generate the group data in B-F. As indicated in A, we carried out intranasal (*I.N.*) instillation of siRNA that was non-targeting (*siNT*) or against presenilin1 (*siPS1*), then intranasally-instilled SA^GFP^, then quantified mouse breathing score (B), lung wet weight to body weight ratio (*LW/BW*) (C), bronchoalveolar lavage (*BAL*) fluid content of protein (D) and leukocytes (E), and lung content of viable SA^GFP^ (F). In B, *n* is indicated in the panel legend; *circles* indicate mean ± SEM; \**p* < 0.05 versus siNT-instilled mice by two-tailed *t* test. In C-F, *circles* indicate *n* and each represent one mouse; bars: mean ± SEM; \**p* < 0.05 as indicated by two-tailed *t* test. BAL fluid contents of protein and leukocytes in D-E were quantified using the same fluid specimens. *CFU,* colony-forming units.

Reports propose that Notch cleavage strengthens barrier function in cultured endothelial cells by a three-step mechanism: (1) Notch1 cleavage exposes the Notch transmembrane domain (TMD); (2) TMD exposure stimulates formation of a plasma membrane protein complex that activates Rac1; (3) Rac1 activation drives endothelial junctional protein assembly (31). We considered that Notch cleavage in the alveolar epithelium might drive alveolar barrier repair through a similar mechanism. To test this possibility, we defined the extent to which junctional proteins are lost, then reassemble after SA toxin exposure and whether Notch has a role in the reassembly mechanism (**Figure 8, A-N**). Our findings show alveolar epithelial-like MLE-12 cells formed E-cadherin-containing junctional protein complexes (**Figure 8, B-C**). While PBS had no effect on E-cadherin (**Figure 8D**), purified Hla (Hla^WT^) caused rapid E-cadherin loss (**Figure 8, E and G**). These data show Hla^WT^ caused dissolution of E-cadherin proteins, in line with findings reported by others (18). While E-cadherin spontaneously recovered within 6 h of Hla^WT^ exposure in cells that were not further treated (**Figure 8, F-G**) or were treated with vehicle solution (**Figure 8, H-I**), it failed to recover in cells treated with DAPT (**Figure 8, H and J-K**), indicating that junctional protein recovery after Hla exposure was Notch cleavage-dependent. Immunofluorescence studies of mouse lungs fixed after intranasal instillation of PBS or SA^GFP^ corroborate the cultured cell data, in that E-cadherin in alveoli was lost within hours of SA^GFP^ instillation but rapidly recovered (**Figure 8, L-N**). Together, these findings show that cultured alveolar epithelial-like cells and mouse lung alveoli rapidly recover junctional protein expression after an injury stimulus that induces junctional protein loss. We interpret that Notch cleavage critically mediates the post-injury recovery of barrier protein expression.

**Figure 8.**
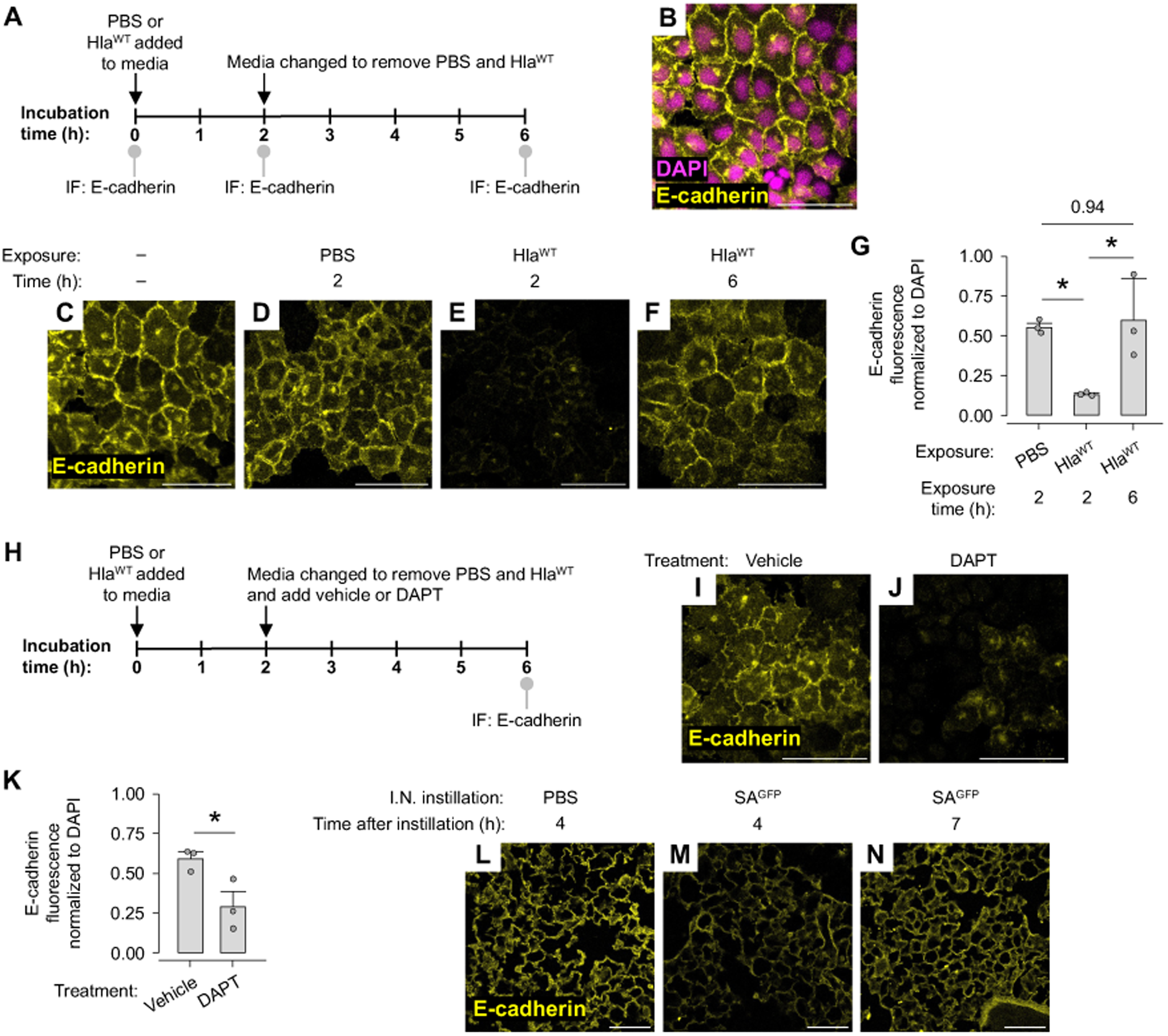
Notch cleavage mediates junctional protein reassembly in alveolar epithelial-like cells. **(A-K)** Cartoons in A and H show the experimental design of the studies used to generate the data shown in B-G and I-K, respectively. As indicated in A and H, we carried out immunofluorescence determinations of E-cadherin expression (*yellow*) by confocal microscopy in cultured alveolar epithelial-like MLE-12 cells before (B-C) and after (D-F and I-J) cell exposure to PBS, purified Hla (*Hla^WT^*), vehicle, and the gamma-secretase inhibitor, DAPT as indicated. Note, Hla^WT^ caused rapid loss of E-cadherin from cell-cell junctions. Junctional E-cadherin spontaneously recovered, but DAPT disrupted the recovery. In the group data in G and K, *circles* indicate *n* and each represent one biological replicate; bars: mean ± SEM; \**p* < 0.05 or as indicated by ANOVA with post hoc Tukey testing (G) and two-tailed *t* test (K). Scale bars: 50 μm. **(L-N)** Confocal images show immunofluorescence of E-cadherin (*yellow*) in tissue sections of mouse lungs that were fixed at the indicated times after intranasal (*I.N.*) instillation of PBS or SA^GFP^ as indicated. Images were replicated 2x per group. Scale bars: 100 μm.

### Therapeutic potential of Notch cleavage to accelerate alveolar barrier repair in lung infection

Approaches that leverage the Notch-mediated barrier repair pathway might be therapeutic in SA lung infection. To test this possibility, we pretreated mice with intranasal instillation of a PS1-encoding plasmid (PS1-OE) (40) to induce PS1 overexpression in the alveolar epithelium (**Figure 9A**). Confocal images show PS1-OE pretreatment increased DsRed-reporter fluorescence in the alveolar epithelium of SA^GFP^-infected lungs (**Figure 9, B-D**), indicating that the PS1-OE plasmid functioned as expected to augment SA^GFP^-induced Notch cleavage in the alveolar epithelium. In mouse models of SA^GFP^ infection, PS1-OE pretreatment did not alter LW/BW ratio or BAL protein content at 6 h after SA^GFP^ instillation (**Figure 9, E-G**), indicating that augmenting Notch cleavage in the alveolar epithelium had no effect on the extent of the initial SA^GFP^-induced lung injury. However, PS1-OE pretreatment reduced LW/BW ratio and BAL protein content at 24 h after SA^GFP^ instillation (**Figure 9, E-G**), indicating that enhancing Notch cleavage in the alveolar epithelium accelerated lung injury resolution. Follow-up experiments show PS1-OE pretreatment did not impact BAL leukocyte content or lung SA^GFP^ content at either 6 h or 24 h after SA^GFP^ instillation (**Figure 9, H-I**), ruling out the possibility that the pro-repair effect of PS1-OE pretreatment stemmed from changes in lung inflammation or bacterial burden. We conclude from these findings that PS1-OE pretreatment augmented SA^GFP^-induced Notch cleavage in the alveolar epithelium, and the augmented Notch cleavage accelerated lung injury resolution after SA^GFP^ lung infection in an inflammation-independent manner. These data strengthen the evidence that Notch cleavage in the alveolar epithelium determines alveolar barrier repair, and they suggest approaches that enhance Notch-mediated barrier-strengthening pathways in alveoli may be therapeutic in SA^GFP^ lung infection.

**Figure 9.**
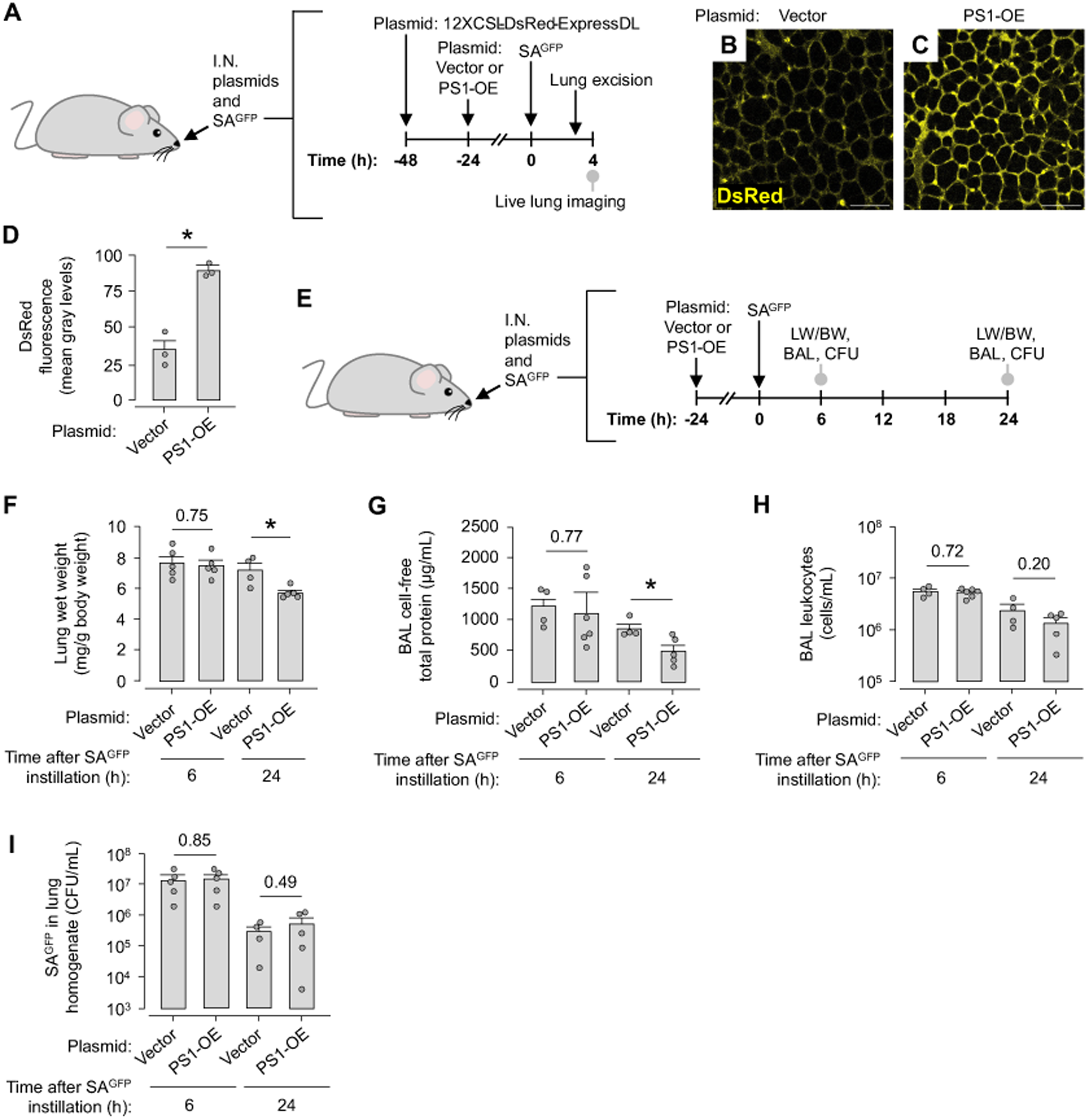
Augmenting Notch cleavage in the alveolar epithelium accelerates lung repair after *S. aureus-*induced acute lung injury. **(A-D)** The cartoon in A shows the experimental design of the studies used to generate the data shown in B-D. As indicated in A, we gave sequential intranasal (*I.N.*) instillations of the 12XCSL-DsRed-ExpressDL plasmid, either vector or presenilin1-overexpression (*PS1-OE*) plasmid, and SA^GFP^, then excised the lungs and viewed plasmid-associated DsRed fluorescence (*yellow*) in live alveoli by confocal microscopy (B-C). In the group data in D, *circles* indicate *n* and each represent one mouse in which mean DsRed fluorescence was quantified in imaging fields of at least 50 alveoli; bars: mean ± SEM; \**p* < 0.05 or as indicated by two-tailed *t* test. Scale bars: 100 μm. **(E-I)** The cartoon in E shows the experimental design of studies used to generate the group data in F-I. As indicated in E, we carried out sequential I.N. instillations of vector or PS1-OE plasmid and SA^GFP^, then quantified lung wet weight to body weight ratio (*LW/BW*) (F), bronchoalveolar lavage (*BAL*) fluid content of protein (G) and leukocytes (H), and lung content of viable SA^GFP^ (I). In F-I, *circles* indicate *n* and each represent one mouse; bars: mean ± SEM; \**p* < 0.05 as indicated by two-tailed *t* test. BAL fluid contents of protein and leukocytes in G-H were quantified using the same fluid specimens. *CFU,* colony-forming units.

A potential problem of leveraging Notch-mediated pathways for barrier-strengthening therapy relates to the NICD. NICD is released when gamma-secretase cleaves Notch to expose the TMD. Subsequently, NICD moves to the cell nucleus and initiates transcriptional events that can associate with lung disease (41), raising the possibility that NICD released in the course of therapeutic exposure of the endogenous Notch TMD could cause harm. To address this issue, we adapted an existing approach (31) to stimulate TMD-mediated barrier repair without inducing NICD release. Thus, we generated plasmids that encode either a Notch1 TMD-GFP fusion protein (TMD-GFP) or GFP under control of a CMV promoter.

Our initial studies aimed to define the effects of plasmid transfection in alveolar epithelial-like cells. Immunofluorescence data show GFP fluorescence of TMD-GFP-transfected cells had a cytosolic distribution in some cells (**Figure 10, A-B**) but colocalized with E-cadherin at plasma membranes in others (**Figure 10, C-D**), indicating that plasmid-derived TMD protein can assume the plasma membrane localization of endogenous TMD. Immunoblot data show TMD-GFP transfection had no effect on Hes1 expression in cultured alveolar epithelial-like cells that were untreated or exposed to Hla^WT^ (**Figure 10E**), indicating that plasmid-induced TMD-GFP expression did not stimulate NICD-mediated transcriptional responses that can associate with Notch-related lung disease.

**Figure 10.**
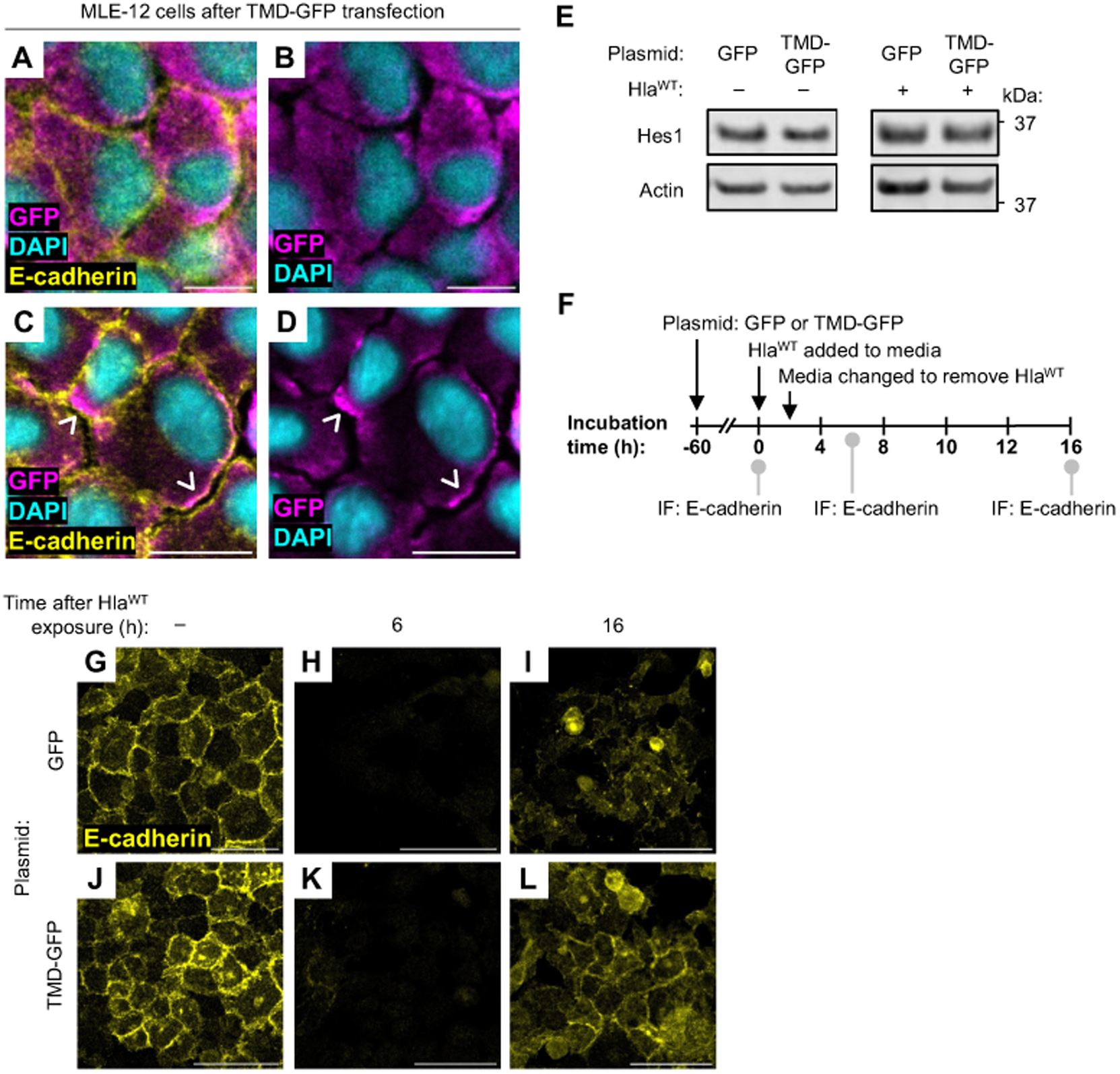
Notch TMD expression promotes junctional protein recovery after Hla-induced junctional protein loss. **(A-D)** Confocal images show immunofluorescence of GFP (*magenta*), DAPI (*cyan*), and E-cadherin (*yellow*) in cultured alveolar epithelial-like MLE-12 cells transfected with plasmid DNA encoding a Notch TMD-GFP fusion protein (*TMD-GFP*). *Arrowheads* in C-D point out locations where GFP and E-cadherin fluorescences merge, signaling TMD-GFP protein localizes to plasma membranes. Scale bars: 20 μm. **(E)** Immunoblot data show Hes1 and actin protein content in MLE-12 cells transfected with plasmid DNA encoding GFP or TMD-GFP, then either untreated or exposed to purified Hla (*Hla^WT^*) as indicated. Hla^WT^ exposure was for 10 min, then the cells were incubated in culture media for 45 min prior to immunoblot processing. Immunoblots were each replicated 2x. **(F-L)** The cartoon in F shows the experimental design of the studies used to generate the data shown in G-L. As indicated in F, we carried out immunofluorescence determinations of E-cadherin expression (*yellow*) by confocal microscopy in MLE-12 cells transfected with plasmid DNA encoding GFP or TMD-GFP. Transfected cells were exposed to Hla^WT^ as indicated. Note, while cells in both groups show a cell junctional pattern of E-cadherin under baseline conditions that is lost in response to Hla^WT^ exposure, only TMD-GFP-transfected cells show E-cadherin recovery. Fluorescence of recovered E-cadherin in GFP-transfected cells appears disorganized and is not located at cell-cell junctions. Images each replicated 2x. Scale bars: 50 μm.

Since Notch1 TMD expression in cultured endothelial cells promotes VE-cadherin assembly (31), we tested whether TMD-GFP transfection in alveolar epithelial-like cells enhances junctional protein expression (**Figure 10, F-L**). Our findings show that both GFP- and TMD-GFP-transfected cells formed E-cadherin-containing junctional protein complexes (**Figure 10, G and J**), and both GFP- and TMD-GFP-transfected cells lost E-cadherin after Hla^WT^ exposure (**Figure 10, H and K**). However, while E-cadherin was rapidly regained at cell-cell junctions in TMD-GFP-transfected cells (**Figure 10L**), it appeared disorganized in GFP-transfected cells (**Figure 10I**). These findings are consistent with the notion that TMD-GFP expression expedites junctional protein recovery after Hla-induced cellular injury.

Finally, we tested the efficacy of TMD-GFP transfection to accelerate barrier repair in mouse models of SA lung infection (**Figure 11, A-F**). Intranasal instillation of each plasmid led to GFP expression in the alveolar epithelium (**Figure 11, A-C**), indicating the plasmids were taken up and expressed by the alveolar epithelium, as expected. Within the first 6 h after intranasal SA^GFP^ instillation, TMD-GFP- and GFP-pretreated mice had similar breathing scores and LW/BW ratios (**Figure 11, D-F**), indicating that TMD-GFP expression had no effect on the extent of the initial lung injury after SA^GFP^ infection. However, TMD-GFP-pretreated mice had lower breathing scores and LW/BW ratio at 24 h after SA^GFP^ instillation (**Figure 11, E-F**), indicating that mice pretreated with TMD-GFP transfection had faster lung injury resolution. These findings show TMD-GFP expression in the alveolar epithelium accelerated lung repair and recovery of barrier function after SA^GFP^-induced lung injury. Taking the mouse and cultured cell data together, we conclude that alveolar epithelial expression of TMD-GFP promoted recovery of junctional proteins and alveolar barrier function to accelerate lung repair after SA lung infection (**Figure 12**).

**Figure 11.**
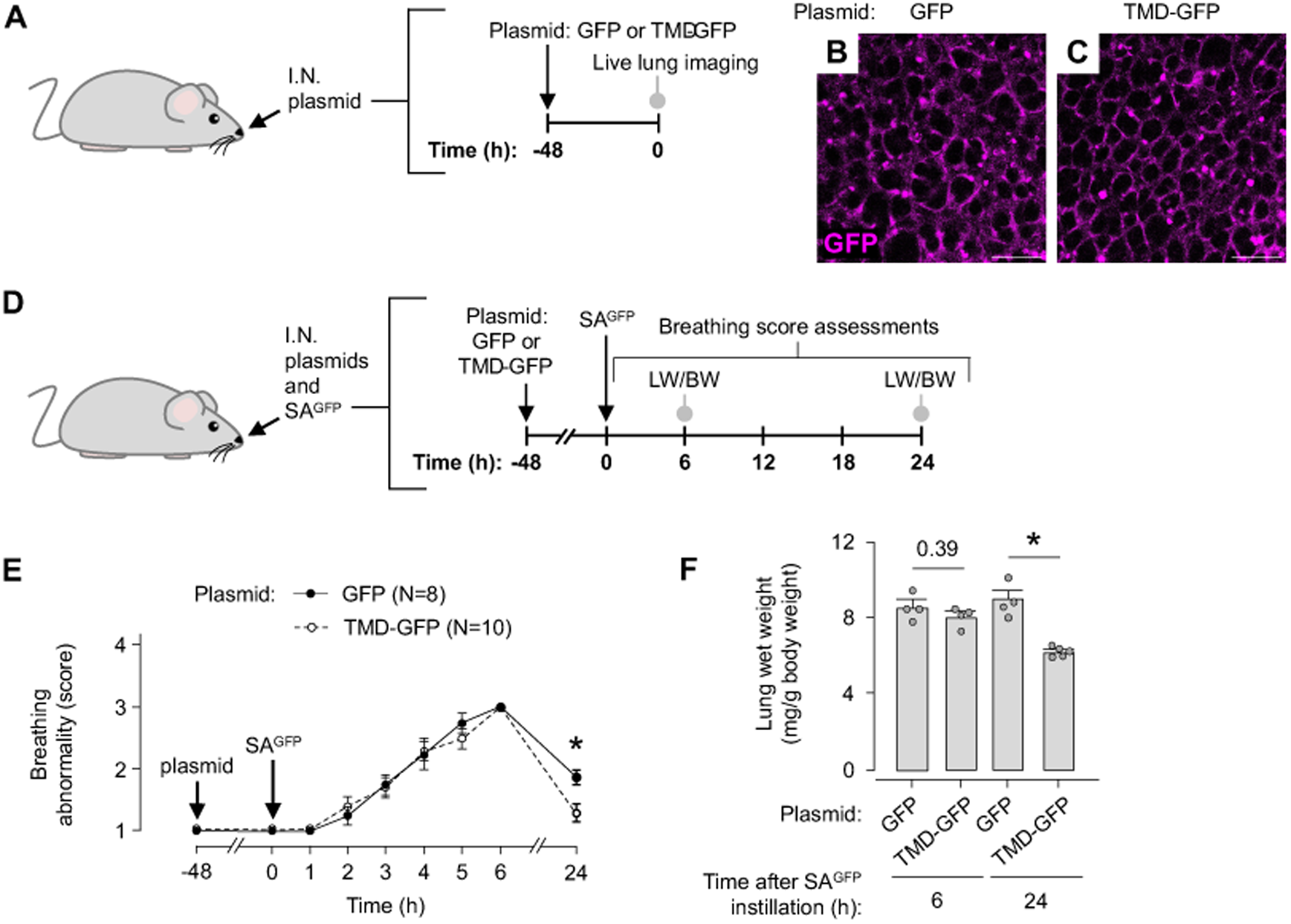
Alveolar epithelial transfection with a Notch TMD-encoding plasmid is therapeutic in mice with *S. aureus-*induced acute lung injury. **(A-C)** The cartoon in A shows the experimental design of the studies used to generate the data shown in B-C. As indicated in A, we gave intranasal (*I.N.*) instillation of plasmid DNA encoding GFP or a Notch TMD-GFP fusion protein (*TMD-GFP*), then excised the lungs and viewed GFP fluorescence in live alveoli (*magenta*) by confocal microscopy (B-C). Scale bars: 100 μm. **(D-F)** The cartoon in D shows the experimental design of the studies used to generate the group data in E-F. As indicated in D, we gave sequential I.N. instillations of the GFP- or TMD-GFP-encoding plasmid and SA^GFP^, then quantified mouse breathing score (E) and lung wet weight to body weight ratio (*LW/BW*) (F). In E, *n* is indicated in the panel legend; *circles* indicate mean ± SEM; \**p* < 0.05 versus TMD-GFP-transfected mice by two-tailed *t* test. In F, *circles* indicate *n* and each represent one mouse; bars: mean ± SEM; \**p* < 0.05 as indicated by two-tailed *t* test.

**Figure 12.**
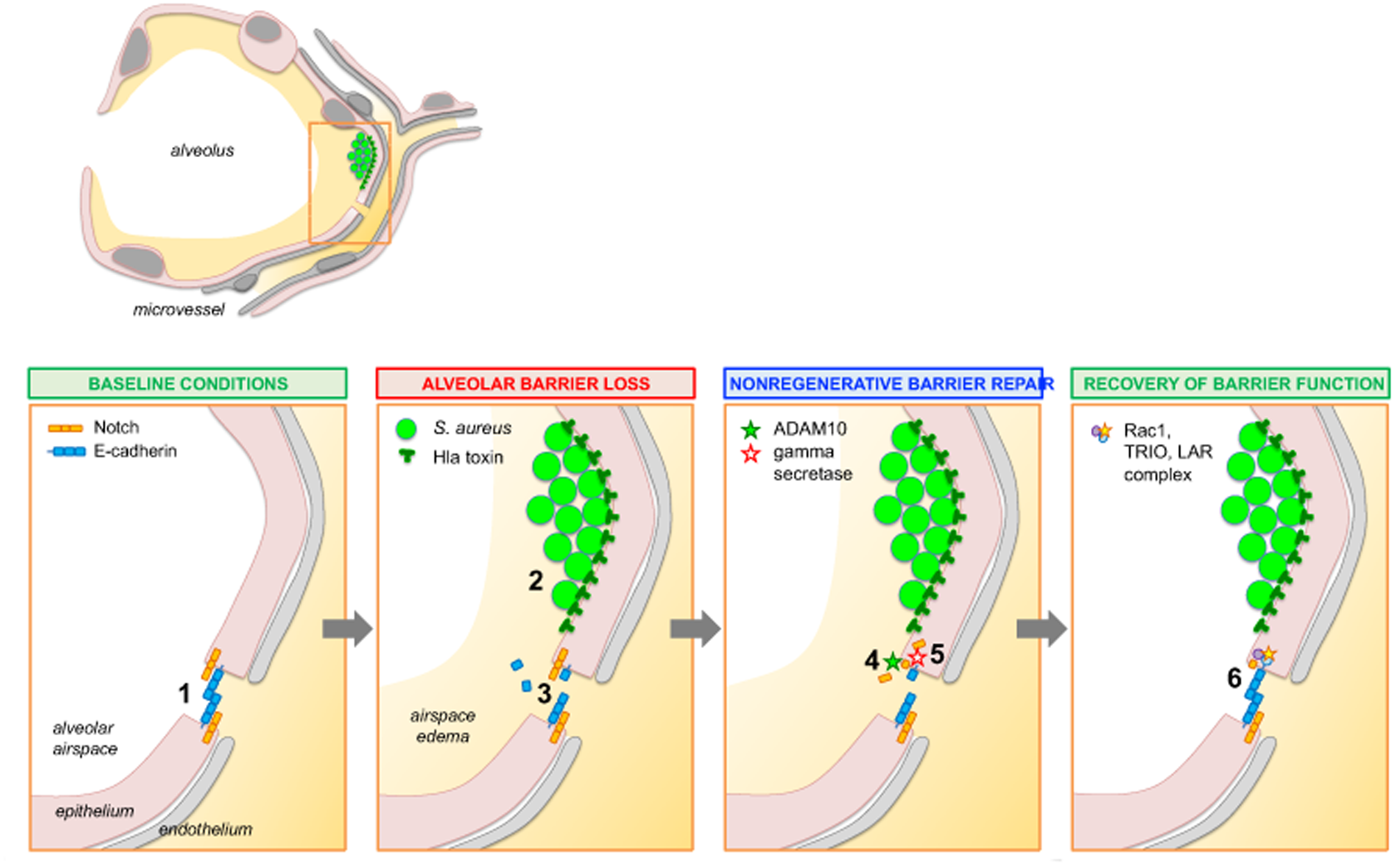
Proposed mechanism of non-regenerative alveolar barrier repair. Cartoons of an alveolus and an alveolar septum (location shown by *dashed rectangle*) illustrate the proposed mechanism. **<u>First panel</u>:** Under baseline conditions, E-cadherin and other intercellular junctional proteins expressed by the alveolar epithelium block the transepithelial transit of liquid from microvascular lumens into airspace lumens, establishing air-blood barrier function **(1)**. **<u>Second panel</u>:** In lung infection, inhaled *S. aureus* form microaggregates in alveolar niches, where they secrete virulence factors such as the pore-forming Hla toxin **(2)**. Hla causes increase of cytosolic Ca^2+^ and ADAM10 activity in the alveolar epithelium, leading to E-cadherin proteolysis and barrier dysfunction. Pulmonary edema forms **(3)**. **<u>Third panel</u>:** Hla and ADAM10 activation in the alveolar epithelium initiates cleavage of Notch at its extracellular, S2 site, leaving behind a Notch fragment consisting of the intracellular and transmembrane domains **(4)**. S2 cleavage stimulates a second Notch cleavage event by gamma-secretase at Notch’s intracellular, S3 site, exposing the Notch transmembrane domain (TMD) **(5)**. **<u>Fourth panel</u>:** TMD exposure catalyzes E-cadherin reassembly, perhaps via formation of a Rac1-containing protein complex, that restores alveolar barrier function **(6)**. Other events that promote lung recovery likely occur concomitantly, including innate immune responses, repair of Hla-mediated epithelial membrane damage, and clearance of airspace edema fluid and protein.

## DISCUSSION

Our findings show, for the first time, that the lung’s air-blood barrier rapidly repairs in a non-regenerative manner after SA-induced acute lung injury. The repair mechanism centers on Notch cleavage in the alveolar epithelium that survives SA-induced alveolar damage. Specifically, we identified that the alveolar epithelium lost barrier function but retained viability in SA-infected lungs. Within hours of barrier loss, barrier function was spontaneously regained through ADAM10- and gamma-secretase-induced Notch cleavage in the alveolar epithelium, which stimulated junctional protein reassembly and recovery of alveolar barrier function. The new understanding gained from our findings is that recovery of alveolar barrier function is a defining feature of lung repair after SA-induced acute lung injury. The alveolar epithelium is thus revealed to be a highly resilient tissue that harbors efficient endogenous mechanisms to stimulate functional recovery after cellular damage. Thus, inhibiting Notch-mediated repair mechanisms in the alveolar epithelium prolongs lung injury in SA-infected lungs, while augmenting them accelerates lung repair.

Although contemporary thinking around lung repair mechanisms focuses on regeneration of the alveolar epithelium that does not survive lung injury (42–44), our findings call attention to pathways by which the alveolar epithelium that survives regains function. We found that Notch cleavage in the alveolar epithelium stimulated the resolution of alveolar barrier hyperpermeability in SA-infected lungs, since inhibition of Notch cleavage in the alveolar epithelium prolonged hyperpermeability and blocked pulmonary edema resolution, while augmentation of Notch cleavage had the opposite effects. These findings build on an established literature that shows other cell types, such as endothelial cells, are similarly capable of regaining barrier function. For example, reports indicate human and rodent endothelial cells spontaneously regain barrier function after exposure to H_2_O_2_ (21) and thrombin (22, 23). In endothelial cells, barrier recovery occurs within minutes of barrier loss, and recovery mechanisms center on junctional protein reassembly (21–23). However, in the published literature there is little understanding of the extent to which the alveolar epithelium recovers barrier function under any conditions. Whether the alveolar epithelium regains barrier function in infected lungs is particularly unclear, since pathogens such as SA cause junctional protein degradation (18), whereas H_2_O_2_ and thrombin cause junctional protein rearrangement (45, 46). Our findings add new understanding of lung repair mechanisms by showing that the alveolar epithelium of SA-infected lungs is, indeed, capable of recovering barrier function. The repair response is surprisingly efficient, since our data show the same stimulus that causes rapid barrier loss – Hla (18, 26) – also stimulated slower, Notch-dependent cellular pathways that led to barrier repair.

A critical aspect of the proposed non-regenerative mechanism of barrier repair is that the alveolar epithelium survives SA-induced lung injury. In this study, we identified that the alveolar epithelium of SA-infected lungs retained viability despite losing barrier function. These findings are supported by work previously published by our group that shows the alveolar epithelium survives Hla-induced plasma membrane pore formation (26), and by work published by others that shows that a considerable proportion of injured epithelial cells survive lung injury by stimuli other than SA and Hla (30, 47). Mechanisms that protect against epithelial cell death in alveoli of SA-infected lungs are poorly understood. Data generated in vitro show protective mechanisms after Hla exposure involve p38 MAP kinase (48), membrane pore closure (49), and membrane pore elimination (50). It is not clear whether these mechanisms protect viability of the alveolar epithelium in situ. Whether they do or not, our findings show that retention of alveolar epithelial viability after SA-induced lung injury paved the way for barrier repair.

Taking our findings together with published data, we propose that the following three-step sequence of events occurs in alveoli of SA-infected lungs (**Figure 12**). The first step is SA-induced alveolar barrier loss. We have shown previously that inhaled SA form microaggregates at alveolar niches, where secreted Hla causes plasma membrane pore formation, cytosolic Ca^2+^ increase, and mitochondrial depolarization in the alveolar epithelium (26). In addition, ADAM10 activation, due either to Hla-ADAM10 interactions (18) or to epithelial cytosolic Ca^2+^ increases (51), leads to E-cadherin degradation (18). The combination of epithelial E-cadherin degradation and mitochondrial depolarization leads to barrier dysfunction in infected alveoli (18, 26). However, barrier dysfunction is not limited to infected alveoli (26). Instead, Ca^2+^ signals are conducted to uninfected alveoli through epithelial gap junctional channels, expanding the scope of damage and leading to widespread airspace edema formation (26). Our findings in this report confirm that SA and Hla induce epithelial junctional protein loss, alveolar barrier dysfunction, and pulmonary edema formation, including in alveoli that are not directly infected.

The second step represents a transition point between alveolar barrier loss and non-regenerative barrier repair (**Figure 12**). Our findings show intranasal SA instillation stimulated Notch cleavage in the alveolar epithelium in a PS1- and ADAM10-dependent manner. A role for PS1 in Notch cleavage in the alveolar epithelium is expected, since PS1 is the catalytic component of the sole Notch-cleaving enzyme, gamma-secretase (36). A role for ADAM10 is also not surprising, since ADAM10 is known to cleave Notch at Notch’s extracellular, S2 cleavage site (37) and thus stimulate PS1-mediated Notch cleavage at Notch’s intracellular, S3 cleavage site (52). However, ADAM10 may not be the only ADAM involved in Notch cleavage in SA lung infection. ADAM17 might also have a role, since ADAM17 also cleaves Notch at its S2 site (53), is expressed by the alveolar epithelium (54), and is implicated in the pathogenesis of SA lung infection (55). Future research might clarify the extent to which ADAM17 contributes to Notch cleavage in alveoli of SA-infected lungs.

The third step is Notch cleavage-induced recovery of alveolar barrier function (**Figure 12**). Reports indicate Hla causes degradation of E-cadherin-containing junctional protein complexes in the alveolar epithelium (18), suggesting that recovery of junctional complexes is a key event in alveolar barrier repair. Our findings confirm that Hla causes E-cadherin degradation and identify that Hla-exposed cells rapidly recovered expression of E-cadherin-containing junctional complexes in a Notch cleavage-dependent manner. These data support the hypothesis that Notch cleavage in the alveolar epithelium of SA-infected lungs stimulates alveolar barrier repair through Notch TMD-mediated junctional complex reassembly. A role for the Notch TMD in barrier function is further supported by reports indicating the Notch TMD strengthens the endothelial barrier in vitro by serving as a scaffold for the formation of a Rac1-containing protein complex that stimulates VE-cadherin assembly at endothelial cell junctions (31). Future studies might identify if Notch-mediated junctional protein reassembly in the alveolar epithelium is also Rac1-dependent, and whether Notch cleavage stimulates assembly of other junctional proteins that determine alveolar barrier function, such as claudins (15, 56, 57).

Our data raise the question of whether interactions between Notch and Notch ligands are required to initiate alveolar barrier repair. Notch cleavage is typically initiated by interactions between Notch and Notch ligands that are expressed either by the Notch-expressing cell (i.e., *cis-*activation of Notch) or an adjacent cell (i.e., *trans-*activation of Notch). The extent to which Notch cleavage in the alveolar epithelium is mediated by Notch-Notch ligand interactions, and whether the interactions are in a *cis* or *trans* configuration, is not yet clear. Published data show deletion of the Notch ligand Dll4 blocks Notch-mediated barrier strengthening in cultured endothelial cells (31), indicating that Dll4 is required for Notch-mediated endothelial barrier strengthening. The alveolar epithelium does not express Dll4 (58), but its expression of other Notch ligands such as Jag1 and Dll1 (32, 41, 59) raises the possibility that Notch-Jag1 or Notch-Dll1 interactions mediate Notch cleavage-induced barrier repair in alveoli.

From a therapeutic perspective, our findings suggest interventions that augment Notch-mediated barrier-strengthening pathways in the alveolar epithelium might represent a new therapeutic approach for SA lung infection. Our data show that two different methods of stimulating Notch-mediated alveolar barrier repair pathways – PS1 overexpression and Notch1 TMD-GFP expression in the alveolar epithelium – each accelerated pulmonary edema resolution after SA infection. Increase of E-cadherin recovery in Notch1 TMD-transfected cells supports the notion that the acceleration of lung repair resulted from recovery of epithelial junctional proteins and regain of alveolar barrier function. Absence of an effect of PS1 overexpression on BAL leukocyte content and lung bacterial burden show that the acceleration was not mediated by PS1-induced alterations of lung inflammation, immune function, or bacterial killing.

An association between Notch activity and lung disease (41, 60) suggests augmenting Notch-mediated pathways in lung infection could cause harm. Since Notch’s mechanistic role in lung disease seems to result from NICD release and NICD-induced transcriptional responses (41, 60), approaches that augment Notch1 TMD exposure without inducing NICD release have potential to promote barrier-strengthening without inducing potentially injurious transcriptional pathways. Notably, we did not identify increased Hes1 protein, a marker of NICD release, at 48 h after TMD-GFP transfection in cultured cells. Further research is needed to define the long-term consequences of enhancing Notch TMD expression or the expression of other proteins involved in Notch-mediated barrier repair.

Finally, our data lay the groundwork for future studies that address important follow-up questions. First, future research might clarify the extent to which our findings, which were generated primarily in mice, apply to humans. Second, it remains unclear which Notch paralog drives alveolar barrier repair. Our studies focus on Notch1, but Notch2 is also expressed by the alveolar epithelium (32) and could have a role in barrier repair mechanisms. Third, it is not clear how Notch1-mediated alveolar barrier repair occurs in relation to other processes involved in lung recovery from infection, such as alveolar fluid clearance, bacterial killing, efferocytosis, and epithelial regeneration. Follow up experiments might define how temporal and spatial aspects of alveolar barrier repair relate to these other lung recovery processes. In addition, future work might identify whether clinical factors, such as antecedent respiratory viral infection, smoking history, or young or old age, disrupt Notch-mediated alveolar barrier repair, and whether such disruptions contribute to clinical heterogeneity in lung infection outcomes.

In conclusion, our findings show the alveolar epithelium survived SA lung infection and rapidly regained barrier function through a Notch-dependent, non-regenerative mechanism. Interventions that augmented Notch-mediated alveolar barrier repair pathways accelerated recovery from SA-induced lung injury. Together, these findings contribute new understanding of lung repair mechanisms, illuminate alveolar responses to bacteria that stimulate lung function recovery, and suggest a molecular framework for the future development of a first epithelium-directed therapy for acute lung injury.

## METHODS

### Experimental design

Experiments were designed according to ARRIVE guidelines. Experimental units were single mice unless otherwise indicated. Mice used for in vivo studies were allocated to groups in a manner that ensured roughly equal mean mouse weight per group. Groups were mixed within cages, and assessments of breathing abnormalities and need for euthanasia were carried out by an investigator blinded to mouse groups. Outcome measures are indicated in figures and legends.

### Sex as a biological variable

Our study used male mice in order to standardize the experiments by avoiding sex hormone-related effects on acute lung injury severity and Notch signaling (61, 62).

### Animals

Mice were Swiss Webster, purchased from Charles River Laboratories and Taconic Biosciences, 25-40 g, and 6-12 weeks old. We anesthetized mice with inhaled isoflurane (4%) and intraperitoneal injections of ketamine (up to 100 mg/kg) and xylazine (up to 5 mg/kg) for intranasal instillations and surgical procedures. For surgeries, we injected the tail vein of anesthetized mice with heparin (50 units; Mylan), then exsanguinated the mice by cardiac puncture. Additional information regarding intranasal instillations and surgeries is provided in the **Supplemental Methods**.

### Solutions

We purchased Ca^2+^- and Mg^2+^-containing DPBS and Ca^2+^- and Mg^2+^-free PBS from Corning. Isolated lungs were perfused with HEPES-buffered solution of pH 7.4 and osmolality 333 mOsm/L and containing 150 mM Na^+^, 5 mM K^+^, 1 mM Ca^2+^, 1 mM Mg^2+^, 140 mM Cl^-^, 10 mM glucose, 4% dextran (70 kDa; Molecular Probes), and 1% FBS (Gemini Bio-Products). Fluorophores and reagents microinstilled into alveoli were dissolved or suspended in HEPES-buffered solution containing 150 mM Na^+^, 5 mM K^+^, 1 mM Ca^2+^, 1 mM Mg^2+^, 140 mM Cl^-^, and 10 mM glucose.

### Reagents

Reagents were freshly constituted for experiments. Hla was purchased from Sigma-Aldrich and reconstituted in deionized water at 0.5 mg/ml, then diluted with DMEM/F-12 medium (catalog 30-2006, ATCC) to 0.01 μg/ml for immunoblot experiments and 1 μg/ml for immunofluorescence experiments. DAPT was purchased from Santa Cruz Biotechnology and reconstituted in DMSO (Fisher Scientific), then diluted to 50 nM with HEPES-buffered solution or DMEM for live lung imaging and cell culture experiments, respectively. CuSO_4_ for fluorescence quenching experiments was prepared from Copper(II) sulfate (Sigma-Aldrich), reconstituted in deionized water, and added to the HEPES-buffered lung perfusate solution or the HEPES-buffered alveolar microinstillation solution at a final concentration of 250 μM CuSO_4_. Saponin (EMD Millipore) at 30 μM was added to the CuSO_4_-containing solutions to facilitate copper entry into live cells without inducing cell death (63). Saponin at 1% w/v was used for alveolar epithelial viability studies to induce rapid cell death (29).

### Fluorophores

We purchased fluorescein isothiocyanate (FITC)-conjugated dextran (20 kDa; 5 mg/mL), tetramethylrhodamine isothiocyanate (TRITC)-conjugated dextran (70 kDa; 10mg/mL), and calcein red-orange AM (10 μM) from ThermoFisher Scientific.

### Bacterial strain, preparation, and inoculation

*S. aureus* was GFP-tagged strain USA300 LAC (SA^GFP^) and GFP-tagged mutant USA300 LAC that is Hla-deficient (SA*^hla::^*^Erm^). Both strains carry chloramphenicol resistance genes. Single bacterial colonies were propagated in autoclaved LB media containing chloramphenicol (10 μg/mL) in a shaking incubator for 18 h (stationary phase) or to OD_600nm_=1 (exponential phase), then intranasally-instilled to deliver 7 x 10^7^ (exponential phase) or 1 x 10^8^ (stationary phase) CFU per mouse. Additional information is provided in the **Supplemental Methods**.

### Isolated, blood-perfused lungs

Using our reported methods (26, 27), we cannulated the trachea, pulmonary artery, and left atrium of the heart of exsanguinated mice, then excised the heart, lungs, and cannulas en bloc for live lung imaging by confocal microscopy. Airway, pulmonary artery, and left atrium pressures were maintained at 6, 10, and 3 cm H_2_O, respectively. Additional information is provided in the **Supplemental Methods**.

### Alveolar microinstillation

We hand-beveled glass micropipettes (Sutter Instruments) to micropuncture single alveoli under bright-field microscopy, as we have done previously (26, 27). Micropunctured alveoli were instilled with reagents in solution, resulting in their spread from the micropunctured alveolus to neighboring alveoli. Microinstillations were performed in 1-3 alveoli bordering each imaging field.

### Live lung imaging and analysis

Using our established methods (26, 27), we viewed alveoli by confocal microscopy (LSM800; Zeiss) with a 20x water immersion objective (NA 1.0; Zeiss) and coverslip. We used bright-field microscopy to randomly select regions of 30-50 alveoli for imaging. Additional information is provided in **Supplemental Methods**.

### Alveolar permeability determinations

We determined alveolar barrier function using our established method (26, 27), in which we transiently add FITC-conjugated dextran (20 kDa) or TRITC-conjugated dextran (70 kDa) to the HEPES-buffered lung perfusate solution described in “Solutions”. Alveolar barrier function was assessed using images generated within 30 min of the dextran addition. To make these assessments, we determined whether each alveolus in each imaging field contained dextran in the airspace. Alveoli were considered dextran-positive if dextran was located within the alveolar airspace and at least 25 gray levels.

### Survival assessments

An investigator blinded to mouse groups assessed need for euthanasia in line with our IACUC-approved protocol. Additional information is provided in the **Supplemental Methods**.

### Breathing score

We used our established method (27) to quantify mouse breathing abnormalities on a 4-point scale based on observed breathing effort (normal or labored), rate (> or < 100 breaths/min), and rhythm (consistent or inconsistent). Higher breathing score correlates with measures of pulmonary edema after acute lung injury (27). Breathing determinations were made by an investigator blinded to mouse groups.

### Lung wet weight to body weight ratio

Body weight was recorded in anesthetized mice at the time of lung excision in untreated mice or at the time of PBS or SA instillation. We used our established methods (26, 27) to exsanguinate anesthetized mice by cardiac puncture and excise and weigh the lungs.

### Protein and leukocyte determinations in BAL fluid

Using our reported methods (26, 27), we cannulated the trachea of exsanguinated mice, then lavaged the lungs with 5 instillations of 1 mL of ice-cold, Ca^2+^-free PBS. For total protein determinations, we centrifuged the first aliquot of BAL fluid return (minimum volume 0.78 mL) for 10 min at 400 *g* and 4°C, then centrifuged the supernatant again for 20 min at 15,000 *g* and 4°C. Total protein was quantified using the Pierce BCA Protein Assay Kit (ThermoFisher). For leukocyte determinations, BAL samples were pooled on a per-mouse basis and centrifuged for 10 min at 500 *g*. The resuspended cells were incubated for 10 min in Turk’s solution (Sigma), then counted using a hemacytometer (Hausser Scientific).

### Bacterial counts

Mouse lungs were mechanically homogenized by crushing in a specimen bag and diluted in 1 mL of DPBS containing Ca^2+^ and Mg^2+^. SA^GFP^ CFU was quantified by serial dilutions on chloramphenicol-containing LB agar plates as we have previously reported (26, 27).

### Preparation, transfection, and instillation of plasmids and siRNA

Plasmids were amplified and purified using an EndoFree Plasmid Maxi Kit (Qiagen). Plasmids included a Notch cleavage reporter (12XCSL-DsRedExpressDL, Addgene #47683); pcDNA3.1 and PS1 in pcDNA3.1 (PS1-OE, gifts of Dr. Alison Goate, Icahn School of Medicine at Mount Sinai); and GFP and Notch1 TMD-GFP each in the pTwist CMV mammalian expression vector (Twist Biosciences). siRNA included SMARTPool ON-TARGETplus siRNA against mouse presenilin1 and mouse non-targeting pool siRNA and was purchased from Dharmacon and reconstituted in molecular grade RNAse-free water (Dharmacon) according to the manufacturer’s instructions. Using our established methods (26, 27), we complexed plasmids with freshy extruded unilamellar liposomes (20 μg/μL; 100 nm pore size; DOTAP; Avanti Lipids) in sterile Opti-MEM (Gibco). For mouse experiments, we administered 75 μg plasmid DNA or 50 μg siRNA per mouse by intranasal instillation. For cell culture experiments, we administered 2.5 μg of plasmid DNA per million cells.

### Immunoblot

Information regarding cell culture procedures is included in the **Supplemental Methods**. MLE-12 cells were incubated with RIPA buffer (ThermoFisher) and Halt protease mix (ThermoFisher) and mechanically removed from cell culture plates using cell scrapers. The resulting solutions were collected in Eppendorf tubes, vortexed every 10 min for 30 min, and centrifuged for 10 min at 15,000 *g* and 4°C. We used the Pierce BCA Protein Assay Kit (ThermoFisher), a plate reader (Molecular Devices) to standardize protein loading in Laemmli 2X Concentrate sample buffer (Sigma), and deionized water for gel electrophoresis (Bio-Rad). Samples were heated to 95°C for 5 min on a heating block (Eppendorf) prior to loading. Band densities were quantified using Image Studio (LI-COR, v.5.2).

### Immunofluorescence

Cultured MLE-12 cells were grown to 100% confluency in 6-well plates, with each well containing one poly-L-lysine-treated cover slip (Neuvitro). Mouse lungs were airway-instilled with OCT, fixed by perfusion and immersion, snap frozen, and sectioned on a cryostat. The cells and lungs were prepared for immunofluorescence studies and imaged as outlined in the **Supplemental Methods**.

### Antibodies

Antibodies were purchased from commercial vendors that provided antibody validation information. Primary antibodies used for immunofluorescence and immunoblot studies included rabbit monoclonal antibody (mAb) against E-cadherin (Cell Signaling; clone 24E10), mouse mAb against GFP (ThermoFisher; clone GF28R), rabbit polyclonal antibody against NICD1 (MilliporeSigma; Anti-Notch1, NT), rabbit mAb against Hes1 (Cell Signaling; clone D6P2U), and rabbit polyclonal antibody against actin (MilliporeSigma). Additional information is provided in the **Supplemental Methods**.

### Statistics

Statistics are indicated in figures and legends. In general, paired comparisons were analyzed using two-tailed *t* tests, multiple comparisons were made using ANOVA with post hoc testing. We considered statistical significance at *p*<0.05. Data were analyzed and figures were prepared using Microsoft Excel and GraphPad Prism (Version 10.3.0).

### Study approval

The Institutional Animal Care and Use Committee of the Icahn School of Medicine at Mount Sinai approved the animal procedures.

## AUTHOR CONTRIBUTIONS

SM and JH designed the study. SM, ST, DC, SS, CS, and JZ contributed to data collection and analysis. All authors contributed to the experimental design and interpretation of results. JH was responsible for the overall project. SM wrote the first draft of the manuscript, and all authors edited the manuscript.

## ACKNOWLEDGEMENTS

This work was supported by NIH grant R01HL164821, Cystic Fibrosis Foundation Research Grant 004792G222, American Lung Association COVID-19 and Emerging Respiratory Viruses Research Award 1031520, and Stony Wold-Herbert Fund, Inc. Research Grant-in-Aid to JH and a Fellowship Grant from the Stony Wold-Herbert Fund, Inc. to SM. We thank Dr. Alison Goate for the PS1-OE plasmid and Dr. Alex Horswill for SA*^hla^*^+^ SA*^hla::^*^Erm^. We thank Ariana Arabadjiev for her contributions to the project’s conceptual design.

## SUPPLEMENTAL METHODS

### Bacterial strain, preparation, and inoculation

Bacteria were stored at -80°C in 25% glycerol in autoclaved Luria-Bertani (LB) broth media (MP Biomedicals) and propagated on LB-agar plates containing chloramphenicol (10 μg/mL) or blood agar plates. Plates were refreshed from frozen stock every 1-2 weeks. For experiments, single bacterial colonies were propagated in autoclaved LB media containing chloramphenicol (10 μg/mL) in a shaking incubator at 37°C and 200 rpm (New Brunswick Scientific) for 18 h (stationary phase) or to OD_600nm_=1 (exponential phase). Stationary phase bacteria were used to generate mouse model data, and exponential phase bacteria were used to generate lung imaging data. Bacteria were prepared for intranasal instillation by centrifuging 1.3 mL of bacterial culture, removing the supernatant, and reconstituting the bacterial pellet in 300 μL of DPBS containing Ca^2+^ and Mg^2+^. Within 40 min of bacterial removal from the incubator, we instilled the bacteria-containing solution by intranasal instillation (30 μL to deliver 7 x 10^7^ or 1 x 10^8^ CFU per mouse for exponential and stationary phase, respectively). For intranasal instillations, mice were rapidly anesthetized and instilled in pairs to ensure similarity of inocula across animals.

### Intranasal instillation

Instillation qualities were recorded on a 4-point scale at the time of instillation by the performing investigator. Quality was considered acceptable for experiments if the instillation was recorded as 3-4 for plasmids and siRNA (i.e., little or no loss of instillate observed) or 4 for bacteria (i.e., no loss of instillate observed).

### Isolated, blood-perfused lungs

Using our established methods (26, 27), we cannulated the trachea, pulmonary artery, and left atrium of the heart of exsanguinated mice, then excised the heart, lungs, and cannulas en bloc. Lungs were inflated with room air through the tracheal cannula and perfused through the pulmonary arterial and left atrial cannulas at 0.4-0.6 mL/min with autologous blood diluted in the HEPES-buffered lung perfusate solution indicated in the “Solutions” section of the Methods. The lung perfusate solution was warmed to 37 C. We used in-line pressure transducers (ADInstruments) to maintain constant airway pressure 6 cm H_2_O via a continuous positive airway pressure machine (Philips Respironics) and pulmonary artery and left atrial pressures 10 and 3 cm H_2_O, respectively, via a roller pump (Ismatec). The lungs were positioned to enable micropuncture and imaging of the diaphragmatic surface of the right caudal lobe or the left lung. Portions of the lung surface that were not used for micropuncture and imaging were covered with plastic wrap to prevent desiccation.

### Live lung imaging and analysis

All images were acquired as single images using Zen (v.2.6; Zeiss) and recorded as Z-sections. Analyzed images were 4-8 μm below the pleura. Optical thickness was 32-34 μm, and frame size was 512 x 512 pixels. We established laser, filter, pinhole, and detector settings at the beginning of each imaging experiment to optimize alveolar fluorescence and avoid fluorescence saturation, then maintained the settings for the duration of the experiment. We confirmed absence of bleed-through between fluorescence emission channels. Images were analyzed using ImageJ (NIH; v.2.0.0-rc-69/1.52n). Brightness and contrast adjustments were applied to individual color channels of entire images and equally to all experiment groups. We did not apply downstream processing or averaging. For determinations of alveolar fluorescence of calcein red-orange, DsRed, or GFP, total intensity across 3 adjacent z-slices was quantified using the Z-project Sum function.

### Survival assessments

Mouse need for euthanasia was determined by a scoring system that included observations of mouse appearance, breathing difficulty, behavior, gait, and response to stimulation by cage top opening and placement on a narrow beam. Mice instilled with SA^GFP^ or PBS were assessed hourly for at least 6 h after instillation, then at least every 12-24 h for up to 3 days. Surviving mice were euthanized at the conclusion of experiments.

### Cell culture

MLE-12 cells were obtained from the American Type Culture Collection (catalog CRL-2110, ATCC) and handled and stored according to the vendor’s instructions. MLE-12 cells were grown as monolayer cultures in DMEM/F-12 medium (catalog 30-2006, ATCC) that was supplemented with 10% FBS (Corning, cat. no. 35-011-CV) and 1% penicillin and streptomycin (Gibco, cat. no. 15140122) and maintained in an incubator at 5% CO_2_ and 37 C.

### Immunofluorescence

Cells were fixed in 4% paraformaldehyde, permeabilized in 0.2% PBST (Tween20: BioRad), and blocked in 5% goat serum (Jackson Laboratories) in PBS without Ca^2+^ or Mg^2+^ (Corning). Primary antibodies were diluted in 0.2% PBST and incubated for 24 h at 4 C. Secondary antibodies were prepared in 0.2% PBST and incubated for 1 h at room temperature. Cells were treated with DAPI 1 μg/ml (Sigma) for 5 min for nuclear staining. Coverslips were removed from the wells with forceps, inverted onto slides containing VectaShield antifade mounting medium (Vector Laboratories, cat. no. H-1000-10), sealed with nail polish, and cured overnight. Mouse lungs were excised, given intratracheal instillation of 0.7 mL OCT (Sakura Finetek), and perfused sequentially with ice-cold PBS and 4% paraformaldehyde (Electron Microscopy Sciences). The trachea and great vessels were ligated and the lungs were immersed in 4% paraformaldehyde for 24 h at 4 C, then dehydrated in sucrose solution (25% then 12.5%) for 48 h at each concentration. The lungs were then embedded in OCT, snap frozen on dry ice, stored at -80C, and sectioned on a Cryostat (Leica CM 3050 S) to generate 10 μm slices of tissue on microscope slides (Fisherbrand). The tissue slices were permeabilized with 0.1% Triton X-100 (ThermoScientific) and blocked with 10% donkey serum (Jackson Laboratories) in 0.05% PBST. Primary antibodies were diluted in 0.05% PBST and incubated for 24 h at 4 C, and secondary antibodies were prepared in 0.05% PBST and incubated for 1 h at room temperature. Tissue was mounted in VectaShield antifade mounting medium with DAPI (Vector Laboratories) and covered with coverslips.

### Antibodies

Primary antibodies used for immunofluorescence studies included rabbit monoclonal antibody (mAb) against E-cadherin (Cell Signaling; clone 24E10; 1:1600 dilution; cat. no. 3195; lot 15) and mouse mAb against GFP (ThermoFisher; clone GF28R; 1:1000 dilution; cat. no. MA5-15256, lot ZI398519). Secondary antibodies were purchased from Jackson Laboratories and included AlexaFluor 488 goat anti-rabbit (1:300 dilution; cat. no. 111-545-003; lot 167908) and AlexaFluor 647 goat anti-mouse (1:300 dilution; cat. no. 115-605-003, lot 141733). Antibodies used for immunoblot studies included rabbit polyclonal antibody against NICD1 (MilliporeSigma; Anti-Notch1, NT; 1:500 dilution; cat. no. 07-1232, lot 3899517), rabbit mAb against Hes1 (Cell Signaling; clone D6P2U; 1:1000 dilution; cat. no. 11988, lot 4), and rabbit polyclonal antibody against actin (MilliporeSigma; 1:2000 dilution; cat. no. A2066, lot 202986). Secondary antibodies were purchased from LI-COR and included IRDye 800CW goat anti-rabbit (cat. no. 925-32211, lot D10629-11) and IRDye 680LT goat anti-rabbit (cat. no. 925-68021). Primary antibodies were diluted in StartingBlock T20 Blocking Buffer (ThermoFisher) and incubated with membranes for 24 h at 4 C (NICD1 and Hes1) or 1 h at room temperature (actin). Secondary antibodies were diluted to 1:10,000 and incubated for 40 minutes at room temperature.

### Immunofluorescence imaging

Samples were imaged by confocal microscopy (LSM800; Zeiss) with a 20x water immersion objective (NA 1.0; Zeiss). We used bright-field microscopy to randomly select regions for imaging. All images were acquired as single images using Zen (v.2.6; Zeiss). Microscope settings were optimized at the beginning of each experiment and maintained for all groups. Images were analyzed using ImageJ (NIH; v.2.0.0-rc-69/1.52n). Brightness and contrast adjustments were applied to individual color channels of entire images and equally to all experiment groups. We did not apply downstream processing or averaging.

## SUPPLEMENTAL DATA

**Supplemental Figure 1.**
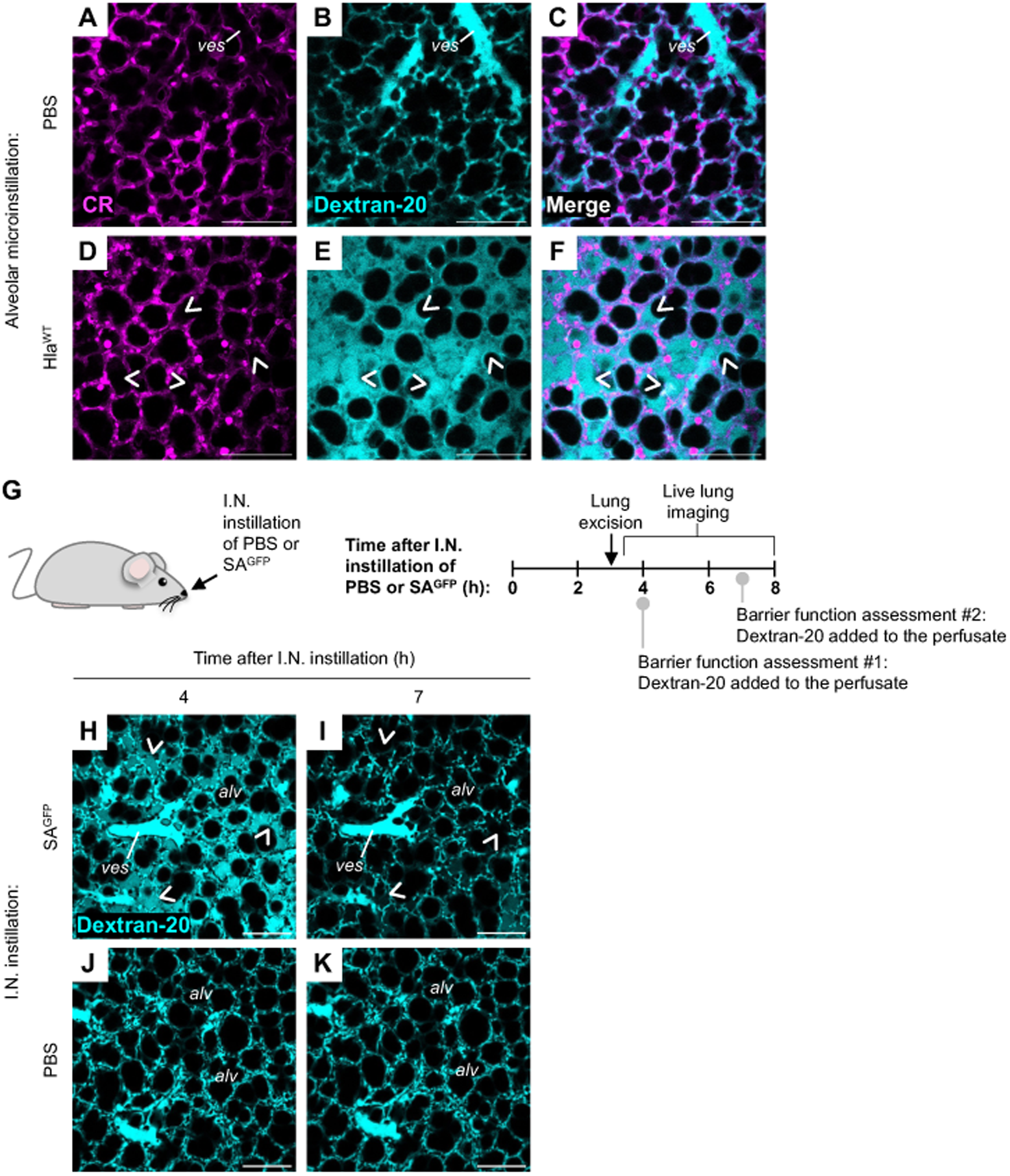
Determinations of alveolar barrier function by live lung imaging. **(A-F)** Confocal images show epithelial fluorescence of calcein red-orange AM (*CR; magenta*) and alveolar fluorescence of FITC-labeled dextran (20 kDa; Dextran-20; *cyan*) in live alveoli of intact, perfused mouse lungs at 4 h after alveolar microinstillation of PBS or purified Hla (*Hla^WT^*) as indicated. Calcein was microinstilled in airspaces by alveolar micropuncture, and dextran was added to the lung perfusate solution. *Arrowheads* indicate Dextran-20 that has leaked into airspaces of Hla^WT^-instilled alveoli in the presence of a dysfunctional alveolar barrier. Note, Dextran-20 is absent from airspaces of PBS-instilled alveoli. *ves,* microvessel. Scale bars: 100 μm. Each set of images was replicated 3 mice. **(G-K)** The cartoon (G) shows the experimental design of studies used to generate the data shown in H-K. As indicated in G, we gave intranasal (*I.N.*) instillation of PBS or SA^GFP^, then excised the lungs and viewed the live alveoli by confocal microscopy (H-K). Alveolar barrier function was assessed at 4 h and again at 7 h after instillation by adding FITC-labeled dextran (20 kDa, *Dextran-20*) to the lung perfusate solution. In H-I, *arrowheads* mark locations where Dextran-20 leaked from microvessels into airspaces at 4 h, but not at 7 h after SA^GFP^ instillation. Note, Dextran-20 is absent from airspaces of the PBS-instilled lungs. Images replicated 2x. Scale bars: 100 μm. *alv;* example alveolus; *ves*, microvessel.

**Supplemental Figure 2.**
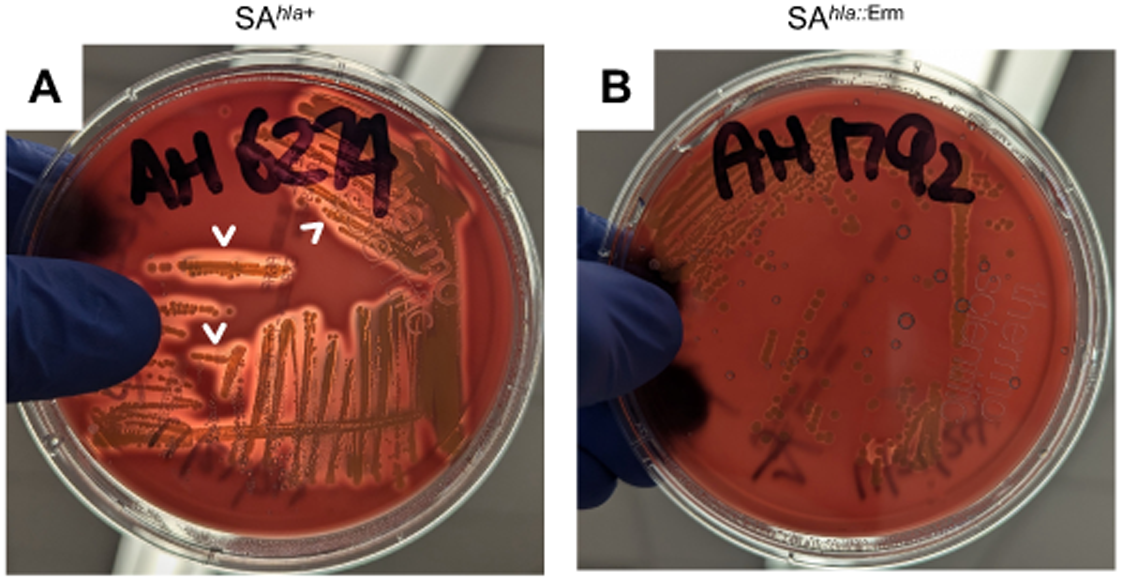
Confirmation of Hla deficiency. **(A-B)** Photos show Hla-producing *S. aureus* USA300 LAC (SA*^hla^*^+^) or Hla-deficient mutant (SA*^hla::^*^Erm^) incubated for 18 h on sheep blood agar. *Arrowheads* indicate zones of hemolytic clearance adjacent to bacterial culture streaks of SA*^hla^*^+^ (A) but not SA*^hla::^*^Erm^ (B). Absence of hemolysis confirms SA*^hla::^*^Erm^ does not secrete Hla.

## REFERENCES

1. Firsova AB, et al. Spatial single-cell atlas reveals regional variations in healthy and diseased human lung. Nat Commun. 2025;16(1):9745.

2. Song L, et al. Cell cross-talk in alveolar microenvironment: from lung injury to fibrosis. Am J Respir Cell Mol Biol. 2024;71(1):30–42.

3. Wagers AJ. The stem cell niche in regenerative medicine. Cell Stem Cell. 2012;10(4):362–9.

4. Sakamoto Y, et al. In-hospital mortality associated with community-acquired pneumonia due to methicillin-resistant *Staphylococcus aureus*: a matched-pair cohort study. BMC Pulm Med. 2021;21(1):345.

5. GBD Antimicrobial Resistance Collaborators. Global mortality associated with 33 bacterial pathogens in 2019: a systematic analysis for the Global Burden of Disease Study 2019. Lancet. 2022;400(10369):2221–48.

6. Rice TW, et al. Critical illness from 2009 pandemic influenza A virus and bacterial coinfection in the United States. Crit Care Med. 2012;40(5):1487–98.

7. McDanel JS, et al. Increased mortality rates associated with *Staphylococcus aureus* and influenza co-infection, Maryland and Iowa, USA. Emerg Infect Dis. 2016;22(7):1253–6.

8. Diep BA, et al. Polymorphonuclear leukocytes mediate *Staphylococcus aureus* Panton-Valentine leukocidin-induced lung inflammation and injury. Proc Natl Acad Sci U S A. 2010;107(12):5587–92.

9. Olsen RJ, et al. Lack of a major role of *Staphylococcus aureus* Panton-Valentine leukocidin in lower respiratory tract infection in nonhuman primates. Am J Pathol. 2010;176(3):1346–54.

10. Weibel ER. Morphometry of the Human Lung. New York, NY: Academic Press Inc.; 1963.

11. Schneeberger-Keeley EE, et al. The ultrastructural basis of alveolar-capillary membrane permeability to peroxidase used as a tracer. J Cell Biol. 1968;37(3):781–93.

12. Schneeberger EE, et al. Substructure of intercellular junctions in freeze-fractured alveolar-capillary membranes of mouse lung. Circ Res. 1976;38(5):404–11.

13. Gorin AB, et al. Differential permeability of endothelial and epithelial barriers to albumin flux. J Appl Physiol Respir Environ Exerc Physiol. 1979;47(6):1315–24.

14. Wiener-Kronish JP, et al. Differential responses of the endothelial and epithelial barriers of the lung in sheep to *Escherichia coli* endotoxin. J Clin Invest. 1991;88(3):864–75.

15. Mitchell LA, et al. Differential effects of claudin-3 and claudin-4 on alveolar epithelial barrier function. Am J Physiol Lung Cell Mol Physiol. 2011;301(1):L40–9.

16. Caraballo JC, et al. Ambient particulate matter affects occludin distribution and increases alveolar transepithelial electrical conductance. Respirology. 2011;16(2):340–9.

17. Bhattacharya J, et al. Regulation and repair of the alveolar-capillary barrier in acute lung injury. Annu Rev Physiol. 2013;75:593–615.

18. Inoshima I, et al. A *Staphylococcus aureus* pore-forming toxin subverts the activity of ADAM10 to cause lethal infection in mice. Nat Med. 2011;17(10):1310–4.

19. Rock JR, et al. Multiple stromal populations contribute to pulmonary fibrosis without evidence for epithelial to mesenchymal transition. Proc Natl Acad Sci U S A. 2011;108(52):E1475–83.

20. Barkauskas CE, et al. Type 2 alveolar cells are stem cells in adult lung. J Clin Invest. 2013;123(7):3025–36.

21. Quadri SK, et al. Resealing of endothelial junctions by focal adhesion kinase. Am J Physiol Lung Cell Mol Physiol. 2007;292(1):L334–42.

22. Birukova AA, et al. Rap-afadin axis in control of Rho signaling and endothelial barrier recovery. Mol Biol Cell. 2013;24(17):2678–88.

23. Aslam M, et al. cAMP controls the restoration of endothelial barrier function after thrombin-induced hyperpermeability via Rac1 activation. Physiol Rep. 2014;2(10).

24. Hidron AI, et al. Emergence of community-acquired methicillin-resistant *Staphylococcus aureus* strain USA300 as a cause of necrotising community-onset pneumonia. Lancet Infect Dis. 2009;9(6):384–92.

25. Carrillo-Marquez MA, et al. *Staphylococcus aureus* pneumonia in children in the era of community-acquired methicillin-resistance at Texas Children’s Hospital. Pediatr Infect Dis J. 2011;30(7):545–50.

26. Hook JL, et al. Disruption of staphylococcal aggregation protects against lethal lung injury. J Clin Invest. 2018;128(3):1074–86.

27. Tang S, et al. Rescue of alveolar wall liquid secretion blocks fatal lung injury by influenza-staphylococcal coinfection. J Clin Invest. 2023;133(19):e163402.

28. Westphalen K, et al. Acid contact in the rodent pulmonary alveolus causes proinflammatory signaling by membrane pore formation. Am J Physiol Lung Cell Mol Physiol. 2012;303(2):L107–16.

29. Manzel LJ, et al. Inhibition by cigarette smoke of nuclear factor-κB-dependent response to bacteria in the airway. Am J Respir Cell Mol Biol. 2011;44(2):155–65.

30. Jansing NL, et al. Unbiased quantitation of alveolar type II to alveolar type I cell transdifferentiation during repair after lung injury in mice. Am J Respir Cell Mol Biol. 2017;57(5):519–26.

31. Polacheck WJ, et al. A non-canonical Notch complex regulates adherens junctions and vascular barrier function. Nature. 2017;552(7684):258–62.

32. Reyfman PA, et al. Single-cell transcriptomic analysis of human lung provides insights into the pathobiology of pulmonary fibrosis. Am J Respir Crit Care Med. 2019;199(12):1517–36.

33. Hernandez SL, et al. *Staphylococcus aureus* alpha toxin activates notch in vascular cells. Angiogenesis. 2019;22(1):197–209.

34. Hansson EM, et al. Recording Notch signaling in real time. Dev Neurosci. 2006;28(1-2):118–27.

35. Islam MN, et al. F-actin scaffold stabilizes lamellar bodies during surfactant secretion. Am J Physiol Lung Cell Mol Physiol. 2014;306(1):L50–L7.

36. Zhao G, et al. Gamma-secretase composed of PS1/Pen2/Aph1a can cleave notch and amyloid precursor protein in the absence of nicastrin. J Neurosci. 2010;30(5):1648–56.

37. van Tetering G, et al. Metalloprotease ADAM10 is required for Notch1 site 2 cleavage. J Biol Chem. 2009;284(45):31018–27.

38. Morohashi Y, et al. C-terminal fragment of presenilin is the molecular target of a dipeptidic gamma-secretase-specific inhibitor DAPT (N-[N-(3,5-difluorophenacetyl)-L-alanyl]-S-phenylglycine t-butyl ester). J Biol Chem. 2006;281(21):14670–6.

39. Roerig DL, et al. First-pass uptake of verapamil, diazepam, and thiopental in the human lung. Anesth Analg. 1989;69(4):461–6.

40. Pimenova AA, et al. Novel presenilin 1 and 2 double knock-out cell line for in vitro validation of PSEN1 and PSEN2 mutations. Neurobiol Dis. 2020;138:104785.

41. Wasnick R, et al. Notch1 induces defective epithelial surfactant processing and pulmonary fibrosis. Am J Respir Crit Care Med. 2023;207(3):283–99.

42. Quantius J, et al. Influenza virus infects epithelial stem/progenitor cells of the distal lung: Impact on Fgfr2b-driven epithelial repair. PLoS Pathog. 2016;12(6):e1005544.

43. McClendon J, et al. Hypoxia-Inducible Factor 1alpha signaling promotes repair of the alveolar epithelium after acute lung injury. Am J Pathol. 2017;187(8):1772–86.

44. Planer JD, et al. After the storm: regeneration, repair, and reestablishment of homeostasis between the alveolar epithelium and innate immune system following viral lung injury. Annu Rev Pathol. 2023;18:337–59.

45. Lee HS, et al. Hydrogen peroxide-induced alterations of tight junction proteins in bovine brain microvascular endothelial cells. Microvasc Res. 2004;68(3):231–8.

46. Gavard J, et al. Protein kinase C-related kinase and ROCK are required for thrombin-induced endothelial cell permeability downstream from Galpha12/13 and Galpha11/q. J Biol Chem. 2008;283(44):29888–96.

47. Hernandez BJ, et al. Intermediary role of lung alveolar type 1 cells in epithelial repair upon sendai virus infection. Am J Respir Cell Mol Biol. 2022;67(3):389–401.

48. Husmann M, et al. Differential role of p38 mitogen activated protein kinase for cellular recovery from attack by pore-forming *S. aureus* alpha-toxin or streptolysin O. Biochem Biophys Res Commun. 2006;344(4):1128–34.

49. Walev I, et al. Recovery of human fibroblasts from attack by the pore-forming alpha-toxin of *Staphylococcus aureus*. Microb Pathog. 1994;17(3):187–201.

50. Husmann M, et al. Elimination of a bacterial pore-forming toxin by sequential endocytosis and exocytosis. FEBS Lett. 2009;583(2):337–44.

51. Le Gall SM, et al. ADAMs 10 and 17 represent differentially regulated components of a general shedding machinery for membrane proteins such as transforming growth factor alpha, L-selectin, and tumor necrosis factor alpha. Mol Biol Cell. 2009;20(6):1785–94.

52. van Tetering G, et al. Proteolytic cleavage of Notch: "HIT and RUN". Curr Mol Med. 2011;11(4):255–69.

53. Brou C, et al. A novel proteolytic cleavage involved in Notch signaling: the role of the disintegrin-metalloprotease TACE. Mol Cell. 2000;5(2):207–16.

54. Dijkstra A, et al. Expression of ADAMs ("a disintegrin and metalloprotease") in the human lung. Virchows Arch. 2009;454(4):441–9.

55. Gomez MI, et al. *Staphylococcus aureus* protein A activates TACE through EGFR-dependent signaling. EMBO J. 2007;26(3):701–9.

56. Li G, et al. Knockout mice reveal key roles for claudin 18 in alveolar barrier properties and fluid homeostasis. Am J Respir Cell Mol Biol. 2014;51(2):210–22.

57. LaFemina MJ, et al. Claudin-18 deficiency results in alveolar barrier dysfunction and impaired alveologenesis in mice. Am J Respir Cell Mol Biol. 2014;51(4):550–8.

58. Xia S, et al. Delta-like 4 is required for pulmonary vascular arborization and alveolarization in the developing lung. JCI Insight. 2021;6(7).

59. Finn J, et al. Dlk1-mediated temporal regulation of Notch signaling is required for differentiation of alveolar type II to type I cells during repair. Cell Rep. 2019;26(11):2942–54 e5.

60. Zhang M, et al. ZEB1-activated LINC01123 accelerates the malignancy in lung adenocarcinoma through NOTCH signaling pathway. Cell Death Dis. 2020;11(11):981.

61. Speyer CL, et al. Regulatory effects of estrogen on acute lung inflammation in mice. Am J Physiol Cell Physiol. 2005;288(4):C881–90.

62. Soares R, et al. Evidence for the Notch signaling pathway on the role of estrogen in angiogenesis. Molecular Endocrinology. 2004;18(9):2333–43.

63. Zou W, et al. Live-cell copper-induced fluorescence quenching of the flavin-binding fluorescent protein CreiLOV. ChemBioChem. 2020;21(9):1356–63.

